# Scalable Enzyme–Reaction Retrieval Enables Metagenomic Biocatalyst Discovery

**DOI:** 10.64898/2026.10.15.759509

**Authors:** Ding Luo, Meiling Jiang, Binju Wang

**Author notes:** These authors contributed equally (Ding Luo, Meiling Jiang). **Corresponding Author Binju Wang** − State Key Laboratory of Physical Chemistry of Solid Surfaces and Fujian Provincial Key Laboratory of Theoretical and Computational Chemistry, College of Chemistry and Chemical Engineering, Xiamen University, Xiamen 361005, P. R. China; Innovation Laboratory for Sciences and Technologies of Energy Materials of Fujian Province (IKKEM), 361005 Xiamen, China.

## Abstract

Enzyme-catalyzed reactions drive metabolism and enable diverse applications in biotechnology and sustainable synthesis, but assigning catalytic functions to uncharacterized enzymes and finding enzymes for target reactions remain major challenges. Despite recent advances in machine learning, translating enzyme function prediction into systematic discovery across the vast space of uncharacterized proteins and biochemical reactions remains challenging. Here we present EzHit, a lightweight framework for bidirectional enzyme–reaction retrieval. Across enzyme annotation, orphan-reaction retrieval and family-specific fine-tuning benchmarks, EzHit achieved strong top-ranked performance and outperformed representative enzyme–reaction matching baselines. We further show that Mahalanobis-distance-based geometric confidence provides more useful reliability estimates than probability-derived uncertainty metrics or evidential learning, enabling confidence-aware prioritization of high-scoring candidates. Owing to its efficiency, EzHit was applied to a nearly trillion-scale OMG_Prot50 × RHEA screen, yielding 8.20 billion retained enzyme–reaction associations across 30.27 million predicted enzyme sequences. Experimental validation further confirmed the activity of EzHit-prioritized metagenomic enzyme candidates for a sparsely annotated reaction. Together, these results establish EzHit as a scalable framework for enzyme annotation, reaction-guided mining and biocatalyst discovery.

## Introduction

Enzymes catalyze the chemical transformations that sustain metabolism and support diverse applications in biotechnology and biomanufacturing.^1–2^ The rapid growth of protein sequence collections and biochemical reaction databases has greatly expanded opportunities for studying enzyme function and discovering new biocatalysts.^3–4^ Sequence databases such as UniProt^5^ and metagenomic protein collections contain vast numbers of protein sequences, many with limited or uncertain functional annotation, while reaction resources such as RHEA^6^, BRENDA^7^ and MetaCyc^8^ catalogue a broad range of biochemical reactions. Despite these growing resources, the catalytic functions of many proteins remain unknown, and suitable enzymes are still difficult to identify for many target reactions. Enzyme function annotation and reaction-guided enzyme mining therefore remain important challenges.

Recent machine learning studies have begun to bridge enzyme sequence space and chemical reaction space. Enzyme Commission (EC) number prediction models such as CLEAN^9^ has improved sequence-level enzyme annotation, whereas enzyme–substrate and enzyme–reaction models such as ESP^10^, CLIPZyme^11^ and EnzymeCAGE^12^ have moved toward reaction-level matching. More recent work has further emphasized this direction. EZSpecificity^13^ uses sequence, structural and substrate information to predict enzyme– substrate specificity, while CATNIP^14^ connects chemical and enzyme sequence space to support enzyme-to-substrate and substrate-to-enzyme prediction in α-ketoglutarate-dependent non-heme iron enzymes. These studies demonstrate the growing potential of machine learning to connect protein sequence information with biochemical function.

Despite this progress, practical challenges remain for enzyme function annotation and reaction-guided enzyme mining. Structural and multimodal methods can provide useful information about how enzymes recognize and catalyze reactions,^12–13^ but they often require reliable protein structures, active-site definitions, or enzyme–substrate poses. These inputs are not always available,^15^ and generating them at scale remains challenging for large protein and reaction databases. More generally, machine-learning models can become overconfident under distribution shift, such that high prediction scores do not necessarily reflect prediction reliability for inputs that differ from the training data.^16^ This issue is particularly relevant to enzyme mining, where candidate proteins and target reactions may extend beyond the distributions represented in curated training datasets. Confidence information can therefore help prioritize high-scoring candidates from large search spaces for experimental testing. However, compared with accuracy-oriented enzyme–reaction retrieval, the use of uncertainty estimates for candidate prioritization remains less systematically explored. This leaves an important gap between benchmark performance and practical candidate selection, where scalability and reliability are both required.

Here we introduce EzHit, a lightweight and confidence-aware framework for enzyme function annotation and reaction-guided enzyme mining. Across enzyme annotation benchmarks and orphan-reaction retrieval tasks, EzHit achieved strong top-ranked performance and outperformed representative baselines across multiple ranking metrics. To improve prediction reliability, we further developed a geometric uncertainty strategy based on Mahalanobis distance for candidate prioritization. The efficiency of EzHit enabled a nearly trillion-pair screen of 30.27 million predicted enzyme sequences from OMG_Prot50^17^ against 30,136 RHEA reactions. After confidence filtering, this screen yielded 8.20 billion retained enzyme–reaction associations and provided ranked candidates for many sparsely annotated reactions. Experimental testing of selected candidates further confirmed that the large-scale screening workflow can recover active enzymes for a sparsely annotated reaction. Together, these results establish EzHit as a scalable framework for enzyme annotation, reaction-guided enzyme mining and biocatalyst discovery.

## Results

### Overview of the EzHit framework and functional applications

To accurately predict the catalytic compatibility between uncharacterized proteins and chemical reactions, we developed EzHit, a comprehensive sequence-based deep learning framework (**Fig. 1a**). By taking amino acid sequences and reaction SMILES strings as direct inputs, the architecture avoids the computational dependency on 3D protein structures. For the protein modality, EzHit leverages the pre-trained protein language model ProtT5^18^ to extract high-dimensional sequence embeddings, capturing complex evolutionary and structural patterns directly from amino acid sequences. For the reaction encoding, we employ a hybrid approach that combines Differential Reaction Fingerprints (DRFP)^19^ and Extended Connectivity Fingerprints (ECFP).^20^ This joint chemical encoding captures both the global reaction mapping rules and local molecular substructures of the reactants and products. Following extraction, the multimodal feature vectors undergo layer normalization and are concatenated to form a unified enzyme-reaction representation. To map this combined feature space to a continuous catalytic probability, we integrated a Kolmogorov-Arnold Network (KAN)^21^ as the prediction head. The KAN module processes these concatenated representations to compute and assign the final predicted catalytic score.

**Figure 1.**
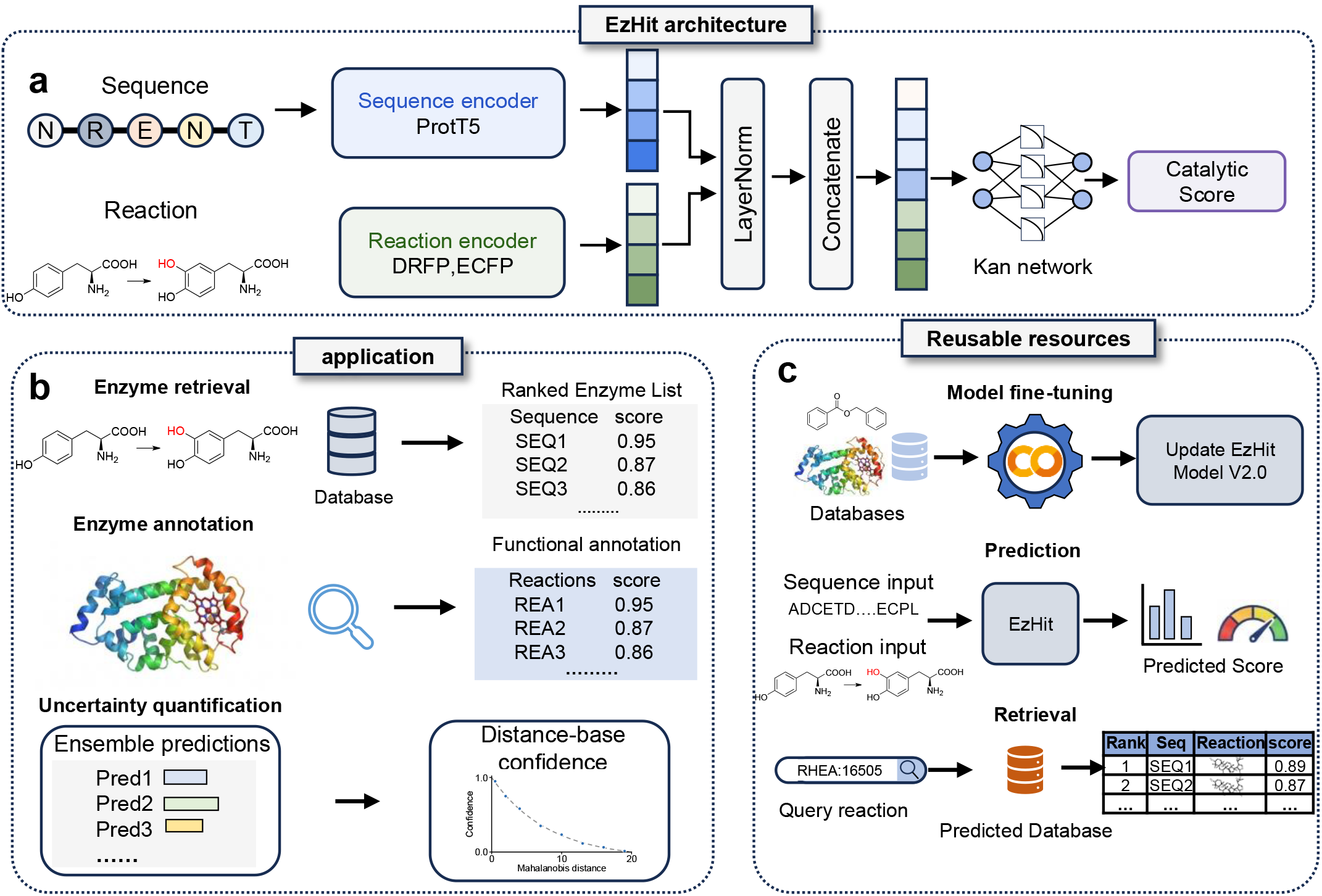
Overview of the EzHit architecture, downstream applications, and reusable prediction resources. **a**, EzHit neural network architecture. Protein sequences are encoded using a ProtT5 sequence encoder, while chemical reactions are represented using Differential Reaction Fingerprint (DRFP) and Extended Connectivity Fingerprint (ECFP) features. The sequence and reaction embeddings are normalized, concatenated, and passed through a Kolmogorov–Arnold Network (KAN) to generate an enzyme–reaction compatibility score for an enzyme– reaction pair. **b**, Representative downstream applications of EzHit. In the reaction-to-enzyme direction, a query reaction can be searched against an enzyme sequence database to retrieve a ranked list of candidate enzymes. In the sequence-to-reaction direction, a query enzyme can be annotated by ranking candidate reactions according to their predicted compatibility. The uncertainty quantification module combines ensemble predictions with Mahalanobis-distance-based confidence estimation, enabling confidence-aware interpretation of predicted enzyme–reaction scores. **c**, Reusable EzHit resources for model adaptation, prediction and retrieval. Custom enzyme–reaction datasets can be used for model fine-tuning, user-defined enzyme–reaction pairs can be scored directly, and precomputed predictions can be queried to retrieve ranked candidates.

To translate enzyme–reaction compatibility scores into practical applications, EzHit supports two complementary downstream tasks (**Fig. 1b**). In reaction-guided enzyme retrieval, a target reaction is used to search an enzyme sequence database and return a ranked list of candidate enzymes according to their predicted enzyme–reaction compatibility. In the reverse direction, EzHit can profile an uncharacterized enzyme against a reaction library and rank candidate reactions for functional annotation. Beyond score-based ranking, EzHit incorporates uncertainty information through ensemble predictions and Mahalanobis-distance-based confidence estimation, allowing candidate pairs to be prioritized by both predicted compatibility and estimated reliability.

To facilitate reuse and broader application, we provide a set of complementary EzHit resources (**Fig. 1c**). Colab notebooks support model fine-tuning on custom enzyme–reaction datasets and direct prediction of user-defined sequence– reaction pairs, while an interactive web interface enables rapid scoring and visualization. For large-scale retrieval, precomputed prediction databases can be queried using standard reaction identifiers such as RHEA IDs to obtain ranked candidate enzymes. Together, these resources support model adaptation, pairwise prediction and database-scale retrieval without requiring users to reproduce the full training and screening pipeline.

### Evaluation of enzyme function prediction on the Enzyme 405 benchmark

To rigorously evaluate the capability of EzHit in predicting functions for uncharacterized enzymes, we benchmarked our model on the recently established Enzyme-405 test set.^12^ This challenging dataset comprises 295 reactions paired with 405 enzymes whose sequences are completely absent from the training data, exhibiting low sequence similarity to known enzymes. The primary objective is to assign each enzyme-reaction pair a confidence score, requiring the model to effectively rank true positive catalytic pairs above a large pool of negative candidates.

We compared EzHit and an ensemble variant (EzHit-Ens) against current state-of-the-art computational baselines: ESP,^10^ CLIPZyme,^11^ and EnzymeCAGE^12^ model. As illustrated in **Fig. 2a–d**, EzHit consistently and significantly outperformed all baseline methods across multiple rigorous ranking and classification metrics. In terms of Success Rate, EzHit achieved approximately 25% at Top-1 and over 65% at Top-10, demonstrating a substantial improvement over EnzymeCAGE (Top-1: *p* < 0.001; Top-10: *p* < 0.01) (**Fig. 2a**). This predictive superiority is further highlighted by the Enrichment Factor (EF) at strict thresholds. EzHit demonstrated an exceptional capacity to prioritize true positive pairs within the top 1% of predictions, achieving an EF of ~45 compared to EnzymeCAGE’s ~33 (*p* < 0.01) (**Fig. 2b**). Correspondingly, EzHit yielded significantly higher scores in Top-10 Discounted Cumulative Gain (DCG) (**Figure. 2c**, *p* < 0.0001) and ROC-AUC (**Fig. 2d**, *p* < 0.01), confirming its robust discriminative power when evaluating structurally novel enzymes. In addition to predictive performance, EzHit showed substantially higher inference throughput than the structure-aware baseline EnzymeCAGE on the Enzyme-405 benchmark, supporting its suitability for large-scale enzyme–reaction screening (**Supplementary Note 6 and Supplementary Table 8**).

**Figure 2.**
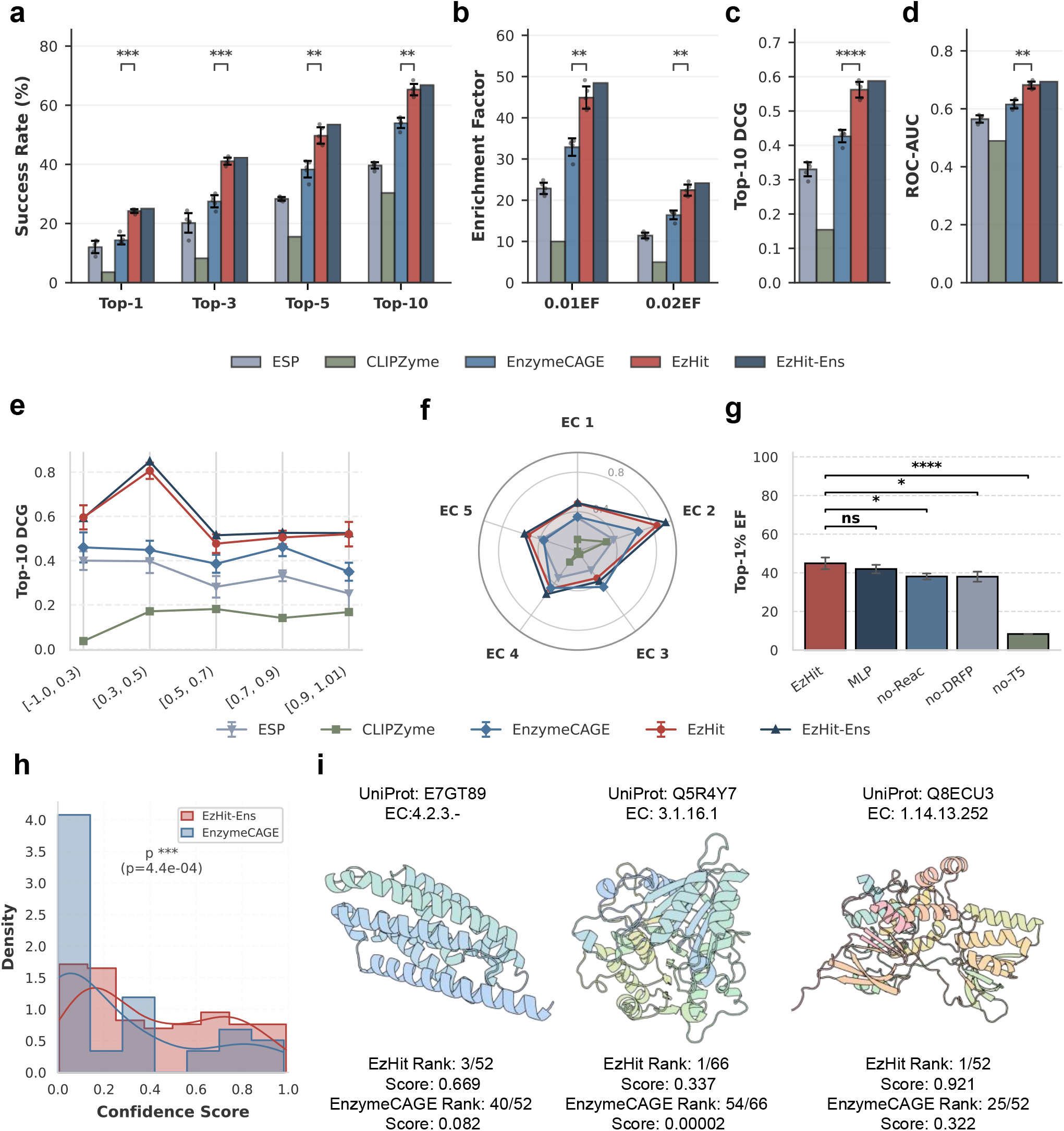
performance evaluation and structural analysis of EzHit on the Enzyme 405 test set. **(a–d)** Comparative evaluation of EzHit against computational baselines (ESP, CLIPZyme, EnzymeCAGE) and an ensemble model. **a**, Performance is measured by Success Rate (%) at Top-1, 3, 5, and 10, **b**, Enrichment Factor at the top 1% and 2% thresholds, **c**, Top-10 Discounted Cumulative Gain (DCG), **d**, and Receiver Operating Characteristic Area Under the Curve (ROC-AUC). **e**, Model performance (Top-10 DCG) evaluated across varying intervals of reaction Morgan fingerprint similarity. **f**, Radar chart illustrating the Top-10 DCG performance across major Enzyme Commission (EC) classes (EC 1 to EC 5). **g**, Ablation study of EzHit evaluated by Top-1% Enrichment Factor. Variants exclude reactant features (no-Reac), differential reaction fingerprints (no-DRFP), or ProtT5 protein language model embeddings (no-T5). **h**, Density distribution of prediction confidence scores for true positive enzyme-reaction pairs predicted by EzHit and EnzymeCAGE. **i**, Structural representations of three representative target enzymes (UniProt IDs: E7GT89, 5R4Y7, 8ECU3) along with their corresponding EC numbers. The individual prediction scores and ranking positions generated by EzHit and EnzymeCAGE are detailed below each 3D structure. For all bar plots (**a–d, g**), data are presented as mean ± s.d. from n = 5 independent runs with different random seeds (40-44). Statistical significance between EzHit and EnzymeCAGE, or between EzHit and the corresponding ablation variants, was assessed using a two-sided paired t-test across matched random seeds: ns, not significant; * *p* < 0.05; ** *p* < 0.01; *** *p* < 0.001; **** *p* < 0.0001. The *p*-value in **h** (*p* = 4.4 * 10^-4^) was calculated using a two-sided Student’s t-test.

To ensure that EzHit’s performance is not heavily biased by reactions highly similar to the training set, we evaluated the Top-10 DCG across varying intervals of reaction Morgan fingerprint similarity (**Fig. 2e**). Consistent trends were also observed across other comprehensive metrics, including Top-k Success Rates and Enrichment Factors at all structural Tanimoto similarity cutoffs (**Supplementary Fig. 2a–f**). EzHit consistently outperformed all baselines across all similarity bins, achieving its highest performance in the [0.3, 0.5) interval, which indicates a robust capacity to generalize to reactions with moderate structural novelty. Furthermore, EzHit demonstrated well-balanced, superior predictive accuracy across diverse major Enzyme Commission (EC) classes (EC 1 through EC 5), yielding particularly notable improvements for Transferases (EC 2) and Hydrolases (EC 3) (**Fig. 2f**).

We then conducted a systematic ablation study to evaluate the contributions of individual feature encodings to the model’s overall success, utilizing the Top-1% EF as the evaluation metric (**Fig. 2g**), supported by extended evaluations on Success Rates and Top-2% EF (**Supplementary Fig. 1a–c**). Replacing the Kolmogorov-Arnold Network (KAN) module with a standard Multi-Layer Perceptron (MLP) resulted in only a minor performance difference on the Enzyme-405 set, indicating that the KAN performs comparably to a standard MLP in this specific scenario. However, this KAN-driven architectural advantage becomes evident when evaluated on the highly sparse and challenging Orphan-335 dataset. While baseline models like EnzymeCAGE show slightly competitive performance at the absolute Top-1 position on sparse data, our KAN-based EzHit architecture demonstrates a more balanced and enriched ranking distribution across all broader ranking metrics (Top-10 SR, EF, and DCG) compared to both the MLP variant and EnzymeCAGE (**Supplementary Fig. 1d–h**). More critically for feature encodings, the absence of either reactant-specific features (no-Reac) or Differential Reaction Fingerprints (no-DRFP) caused significant decrease in performance (no-Reac: *p* < 0.05; no-DRFP: *p* < 0.05, respectively). Furthermore, removing the ProtT5 protein language model embeddings (no-T5) resulted in a substantial drop in predictive accuracy (*p* < 0.0001). These results demonstrate that combining information from the PLM with comprehensive chemical representations is crucial for the model’s robust performance.

To characterize how the models assign confidence to correct predictions, we analyzed the density distribution of scores for true positive enzyme-reaction pairs (**Fig. 2h**). For EnzymeCAGE, the majority of scores for these true positives were clustered in the low-confidence range, reflecting a tendency to assign low scores even to validated functional matches. In contrast, EzHit-Ens yielded a significantly right-shifted distribution toward higher confidence scores (*p* = 4.4 * 10^-4^), demonstrating that our framework provides more robust and discriminative scoring when identifying true catalytic relationships.

To examine these quantitative results through specific biological examples, we analyzed three representative case studies across diverse enzyme classes (**Fig. 2i**). For the lyase E7GT89 (EC 4.2.3.-), EzHit successfully ranked this target enzyme at position 3 out of 52 potential candidates for its known reaction (Score: 0.669), EnzymeCAGE placed it much lower in the ranking (Rank 40/52). Notably, for the nuclease 5R4Y7 (EC 3.1.16.1) and the oxidoreductase 8ECU3 (EC 1.14.13.252), EzHit accurately identified the correct enzymes as the top candidates (Rank 1) with high confidence scores (0.337 and 0.921, respectively). In these instances, the same enzymes were assigned lower priority by EnzymeCAGE, which ranked them at 54/66 and 25/52. These examples illustrate that EzHit can successfully identify active enzymes in scenarios that present difficulties for existing geometric approaches, demonstrating its practical value for the annotation of newly sequenced proteins.

### Accurate enzyme retrieval for orphan reactions

Having evaluated EzHit for enzyme annotation, we next assessed the inverse task of retrieving candidate enzymes for target reactions. This task is particularly relevant for orphan or poorly annotated reactions, where the reaction is known but the corresponding enzyme candidates remain difficult to identify. We used the Orphan-335 benchmark,^12^ in which target reactions are absent from the training set, to evaluate retrieval performance under a reaction-level generalization setting.

EzHit and its ensemble variant, EzHit-Ens, were benchmarked against ESP, CLIPZyme, and EnzymeCAGE using multiple retrieval metrics (**Fig. 3a–d**). At the strict Top-1 cutoff, EnzymeCAGE showed competitive performance, whereas EzHit and EzHit-Ens achieved clearer advantages as the retrieval range expanded. Specifically, EzHit significantly outperformed EnzymeCAGE in Success Rate at Top-10 (Top-10: *p* < 0.05). EzHit-Ens further improved the overall top-k retrieval performance, indicating that the ensemble strategy provided more stable candidate ranking across retrieval cutoffs. These results suggest that EzHit does not only identify the top candidate in favorable cases, but also produces a more informative ranked candidate list for downstream screening.

**Figure 3.**
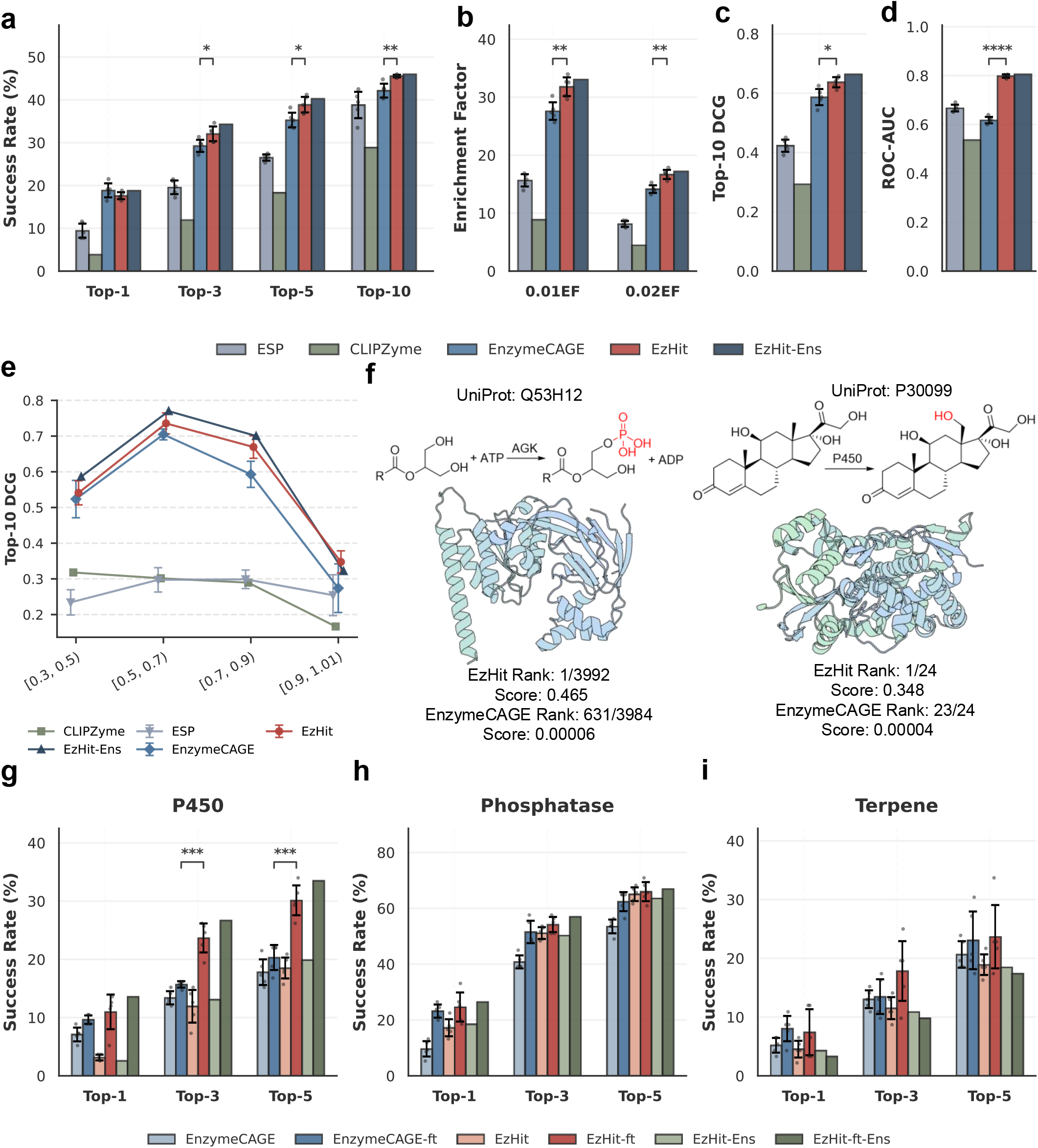
Performance evaluation of EzHit on the Orphan 335 test set and fine tuning efficacy across specific enzyme families. **(a–d)** Comparative evaluation of EzHit against computational baselines (ESP, CLIPZyme, EnzymeCAGE) and an ensemble model (EzHit-Ens) on the Orphan-335 test set. **a**, Performance is measured by Success Rate (%) at Top-1, 3, 5, and 10, **b**, Enrichment Factor at the top 1% and 2% thresholds, **c**, Top-10 Discounted Cumulative Gain (DCG), **d**, and Receiver Operating Characteristic Area Under the Curve (ROC-AUC). **e**, Model performance (Top-10 DCG) evaluated across varying intervals of reaction Morgan fingerprint similarity. **f**, Reaction schemes and structural representations of two representative target enzymes (UniProt IDs: 53H12, P30099). The individual prediction scores and ranking positions generated by EzHit and EnzymeCAGE are detailed below each 3D structure. **(g–i)** Performance evaluation of base models and their fine-tuned counterparts (-ft) across three specific enzyme families: **g**, P450, **h**, Phosphatase, and **i**, Terpene. Performance is measured by Success Rate (%) at Top-1, 3, and 5. For all bar plots (**a–d, g–i**), data are presented as mean ± s.d. from n = 5 independent runs with different random seeds (40-44). Statistical significance between EzHit and EnzymeCAGE (**a–d**), or between base and fine-tuned models (**g–i**), was assessed using a two-sided paired t-test across matched random seeds: ns, not significant; * *p* < 0.05; ** *p* < 0.01; *** *p* < 0.001; **** *p* < 0.0001.

The same trend was observed across additional ranking metrics. EzHit and EzHit-Ens achieved higher Enrichment Factors at the top 1% and 2% of ranked candidates than the baseline methods (**Fig. 3b**; 0.01EF, *p* < 0.05; 0.02EF, *p* < 0.05).

EzHit also improved Top-10 DCG, reflecting better placement of true enzymes within the top-ranked candidate set (**Fig. 3c**; *p* < 0.05). In addition, ROC-AUC analysis showed that EzHit and EzHit-Ens improved the global discrimination between positive and negative enzyme–reaction pairs (**Fig. 3d**; *p* < 0.0001). Together, these metrics indicate that EzHit improves both early retrieval and the overall quality of candidate ranking for orphan reactions.

To further examine whether this advantage depends on reaction similarity to the training set, we stratified Orphan-335 reactions by Morgan fingerprint Tanimoto similarity. EzHit and EzHit-Ens maintained higher Top-10 DCG than the baseline methods across the similarity intervals, including reactions with lower similarity to the training reactions (**Fig. 3e**). This trend indicates that the retrieval performance of EzHit is not limited to orphan reactions that are structurally close to known training reactions. Consistent patterns were also observed in stratified analyses of Top-k Success Rate and Enrichment Factor under different structural similarity cutoffs (**Supplementary Fig. 3a– f**), further supporting the stability of EzHit retrieval across reaction novelty levels.

We next examined two representative retrieval cases from the Orphan-335 benchmark to illustrate how the ranked outputs differ between methods (**Fig. 3f**). In the first case, the target enzyme was an acylglycerol kinase (UniProt: 53H12) associated with an ATP-dependent phosphorylation reaction. Among 3,992 candidate enzymes, EzHit ranked the known positive enzyme at Rank 1 with a score of 0.465. In comparison, EnzymeCAGE ranked the same enzyme at 631/3,984 with a score of 0.00006. In the second case, involving a cytochrome P450-catalyzed hydroxylation reaction, EzHit ranked the target enzyme (UniProt: P30099) at Rank 1 among 24 candidates, with a score of 0.348, whereas EnzymeCAGE ranked the same enzyme at 23/24. These examples illustrate that EzHit can place true target enzymes near the top of large or structurally challenging candidate pools, providing more practical ranked lists for orphan-reaction enzyme retrieval.

### Efficacy of domain specific fine tuning

Enzyme families often differ in substrate scope, catalytic mechanisms, and sequence–function relationships. We therefore asked whether EzHit could be further adapted to enzyme-family-specific retrieval tasks through fine-tuning. We selected three representative enzyme classes with distinct catalytic characteristics: cytochrome P450s,^22^ phosphatases,^23^ and terpene synthases.^24^ For each family, we fine-tuned EzHit and evaluated the resulting single-model and ensemble variants, denoted EzHit-ft and EzHit-ft-Ens, against the corresponding base models and fine-tuned EnzymeCAGE baselines.

Domain-specific fine-tuning improved EzHit performance across the evaluated families (**Fig. 3g–i**). The largest improvement was observed for cytochrome P450s (**Fig. 3g**). Compared with the base EzHit model, EzHit-ft achieved substantially higher Success Rates at broader top-k cutoffs. EzHit-ft also outperformed the fine-tuned EnzymeCAGE baseline at Top-3, Top-5 cutoffs (Top-3, *p* < 0.01; Top-5, *p* < 0.001), suggesting that the EzHit architecture can effectively benefit from family-specific supervision in this chemically specialized enzyme class. For phosphatases, fine-tuning also improved retrieval performance (**Fig. 3h**). EzHit-ft and EzHit-ft-Ens achieved higher or comparable Success Rates relative to the base EzHit model and the fine-tuned EnzymeCAGE variants across Top-5 cutoffs (Top-5, *p* < 0.05). Although the magnitude of improvement was less pronounced than that observed for P450s, the fine-tuned EzHit models consistently remained among the strongest-performing methods. For terpene synthases, the performance gain from fine-tuning was more modest (**Fig. 3i**). Nevertheless, EzHit-ft and EzHit-ft-Ens remained competitive with the strongest baselines across the evaluated top-k cutoffs, and showed improved or comparable Success Rates. This suggests that the benefit of family-specific fine-tuning may vary across enzyme classes, potentially reflecting differences in dataset size, reaction diversity, and family-specific sequence–reaction constraints.

Overall, these results show that EzHit supports both general orphan-reaction retrieval and enzyme-family-specific adaptation. The base model provides strong retrieval performance across structurally diverse orphan reactions, while fine-tuning further improves performance in specialized enzyme families. To facilitate practical application, we further provide a user-friendly Colab workflow that supports EzHit-based prediction, retrieval, and optional fine-tuning on user-defined enzyme-family datasets, enabling broader use of the model without requiring local deployment.

### Geometric ncertainty Quantification Mitigates Out of Distribution Overconfidence

Reliable enzyme annotation requires not only accurate predictions but also the ability to recognize when a prediction is uncertain, especially for out-of-distribution (OOD) enzymes that are poorly represented in the training data. To evaluate whether EzHit can distinguish reliable from unreliable predictions, we systematically examined multiple uncertainty quantification (U) strategies, including probability-based uncertainty metrics derived from ensemble predictions and geometry-based metrics computed from the model’s latent representation (**Fig. 4a**).

**Figure 4.**
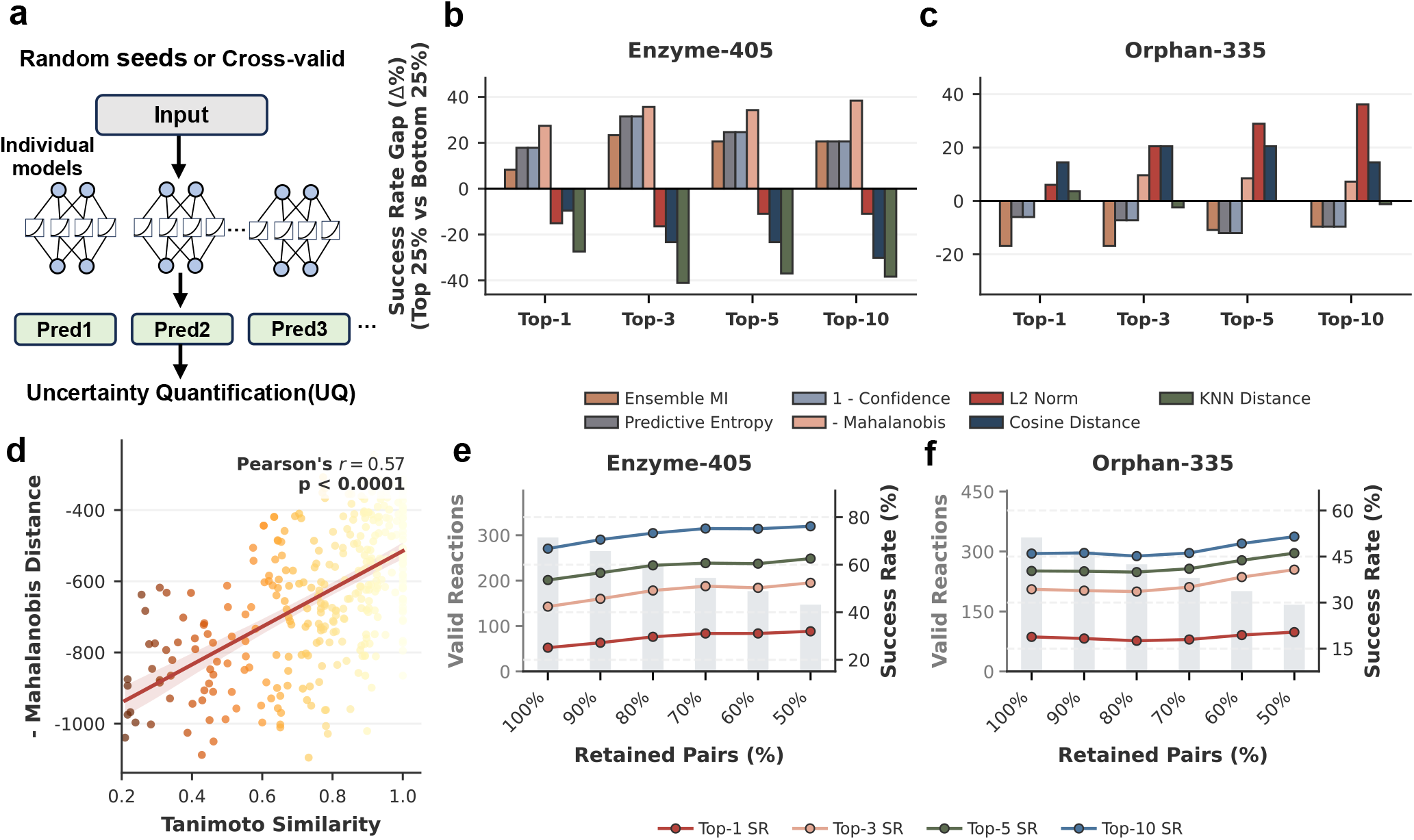
ncertainty uantification and confidence based rejection for reliable enzyme reaction pair prediction. **a**, Schematic overview of the uncertainty quantification (U) pipelines, utilizing both ensemble-based predictions for probability metrics and latent feature space representations for geometric distance metrics. **b, c**, Evaluation of U metric calibration, measured by the Success Rate Gap (Δ%) between the top 25% most confident and bottom 25% least confident predictions. Performance comparisons are shown across traditional probability-based methods (Ensemble MI, Predictive Entropy, 1 - Confidence) and advanced feature-space geometric strategies (-Mahalanobis, L2 Norm, Cosine Distance, KNN Distance) on the Enzyme-405 (**b**) and the out-of-distribution Orphan-335 (**c**) test sets. The inverted Mahalanobis distance (-Mahalanobis) demonstrates uniquely robust calibration across both datasets. **d**, Correlation between the proposed geometric uncertainty (-Mahalanobis Distance) and the maximum structural Tanimoto Similarity to the training set for reactions in the Orphan-335 dataset. The solid line represents the linear regression fit. The positive Pearson’s correlation coefficient (r = 0.57, p < 0.0001) confirms that structurally novel reactions (lower similarity) correctly yield higher uncertainty scores. **e, f**, Confidence-based retention curves illustrating the trade-off between the model Success Rate (line plots, right y-axis) and the number of retained valid reactions (bar plots, left y-axis). Reactions are progressively filtered by retaining the top percentages (from 100% down to 50%) of the most confident predictions, now prioritized based on the - Mahalanobis distance, for the Enzyme-405 (**e**) and Orphan-335 (**f**) datasets. The consistent upward trends validate the metric’s utility in guiding reliable wet-lab experimental designs.

We first assessed whether each U score could stratify predictions by reliability using the success rate gap between the top 25% most confident predictions and the bottom 25% least confident predictions. On the Enzyme-405 dataset, conventional probability-derived metrics, including ensemble mutual information,^25^ predictive entropy, and 1-confidence,^26–27^ showed only limited or inconsistent separation across Top-1 to Top-10 success rates (**Fig. 4b**). In contrast, the inverted Mahalanobis distance^28–29^ produced the most consistent positive success rate gaps across all Top-k metrics, indicating that predictions located closer to the training distribution in latent space are substantially more reliable. This advantage was more pronounced on the highly sparse Orphan-335 dataset, where most probability-based metrics again failed to provide stable discrimination, whereas the Mahalanobis-based score maintained a clear separation between high- and low-confidence predictions (**Fig. 4c**). The maximum success rate gap reached approximately 38 percentage points, demonstrating the ability of geometry-aware uncertainty to identify predictions with markedly different reliability.

We next asked whether this Mahalanobis-based confidence score reflects meaningful distributional proximity rather than merely ranking predictions by model output magnitude. Using Tanimoto similarity as a proxy for reaction-level similarity to known examples, we observed a clear positive correlation between the inverted Mahalanobis distance and Tanimoto similarity (*r* = 0.57, *p* < 0.0001; **Fig. 4d**). Thus, reactions that are more similar to the training distribution tend to occupy more central regions of the learned latent space, whereas structurally or chemically unfamiliar reactions are assigned lower Mahalanobis-based confidence. This result supports the interpretation that the Mahalanobis score captures chemically meaningful novelty in the latent representation.

The practical value of this geometric U strategy was further confirmed by confidence-based retention analysis. As predictions with high Mahalanobis uncertainty were progressively filtered out, the retained reaction pairs exhibited steadily improved Top-k success rates on both Enzyme-405 and Orphan-335 (**Fig. 4e, f**). Importantly, this improvement was achieved while retaining a substantial number of valid reaction pairs, suggesting that Mahalanobis-based filtering can improve experimental prioritization without excessively reducing candidate coverage. By contrast, probability-derived uncertainty metrics produced flat or unstable retention trends, indicating that softmax- or ensemble-probability-based confidence alone provides limited guidance for identifying reliable predictions under OOD conditions (**Supplementary Fig. 4 7**).

To further exclude the possibility that this failure arose from insufficient ensemble diversity, we constructed an additional ensemble using structurally defined cross-validation splits. However, the resulting probability-based U metrics exhibited similar OOD overconfidence behavior and did not recover the reliability separation achieved by Mahalanobis-based scoring (**Supplementary Fig. 8**). We also evaluated evidential deep learning(**Supplementary Note 2, 3**),^30^ which explicitly parameterizes predictive uncertainty through a Dirichlet distribution. Although this approach directly regularizes uncertainty estimation, it substantially reduced the base predictive performance of the KAN-based model across Top-k metrics, suggesting a trade-off between evidential regularization and representation capacity in this setting (**Supplementary Fig. 9**). Together, these analyses indicate that the limitation is not simply due to ensemble construction or the absence of explicit uncertainty regularization, but rather reflects a broader weakness of probability-derived uncertainty for recognizing structurally novel enzyme-reaction pairs.

Overall, these results demonstrate that distribution-aware geometric uncertainty, particularly the inverted Mahalanobis distance in the learned latent space, provides a more reliable indicator of prediction confidence than conventional probability-based U metrics. This enables EzHit to identify lower-confidence predictions for unfamiliar reactions and prioritize candidates that are more likely to succeed in downstream experimental validation.

### Metagenome scale mapping reveals candidate enzymes for underannotated reactions

A major challenge in reaction-guided enzyme mining is that many biochemically defined reactions remain connected to few experimentally characterized protein sequences. We therefore applied EzHit at metagenomic scale to examine whether large-scale enzyme–reaction retrieval could expand the candidate sequence space associated with underannotated chemistry. After filtering OMG_Prot50 with a sequence-level enzyme classifier (**Supplementary Note 5**), we screened 30.27 million predicted enzyme-like proteins against the RHEA reaction database (**Fig. 5a**). Applying a retained-hit threshold of predicted probability > 0.5 and Mahalanobis distance < 20 yielded 8.20 billion protein–reaction associations spanning 19,489 RHEA reactions (**Fig. 5b**). A more stringent threshold of predicted probability ≥ 0.8 and Mahalanobis distance ≤ 16 identified candidate sequences for reactions with limited existing enzyme annotation. Among reactions without linked enzyme annotations in the metadata used for this analysis, 403 received at least one strict candidate, while a further 1,124 reactions with only one or two known enzyme annotations received strict candidates (**Fig. 5c**). These results substantially expand the sequence-level candidate space associated with sparsely annotated RHEA chemistry.

**Figure 5.**
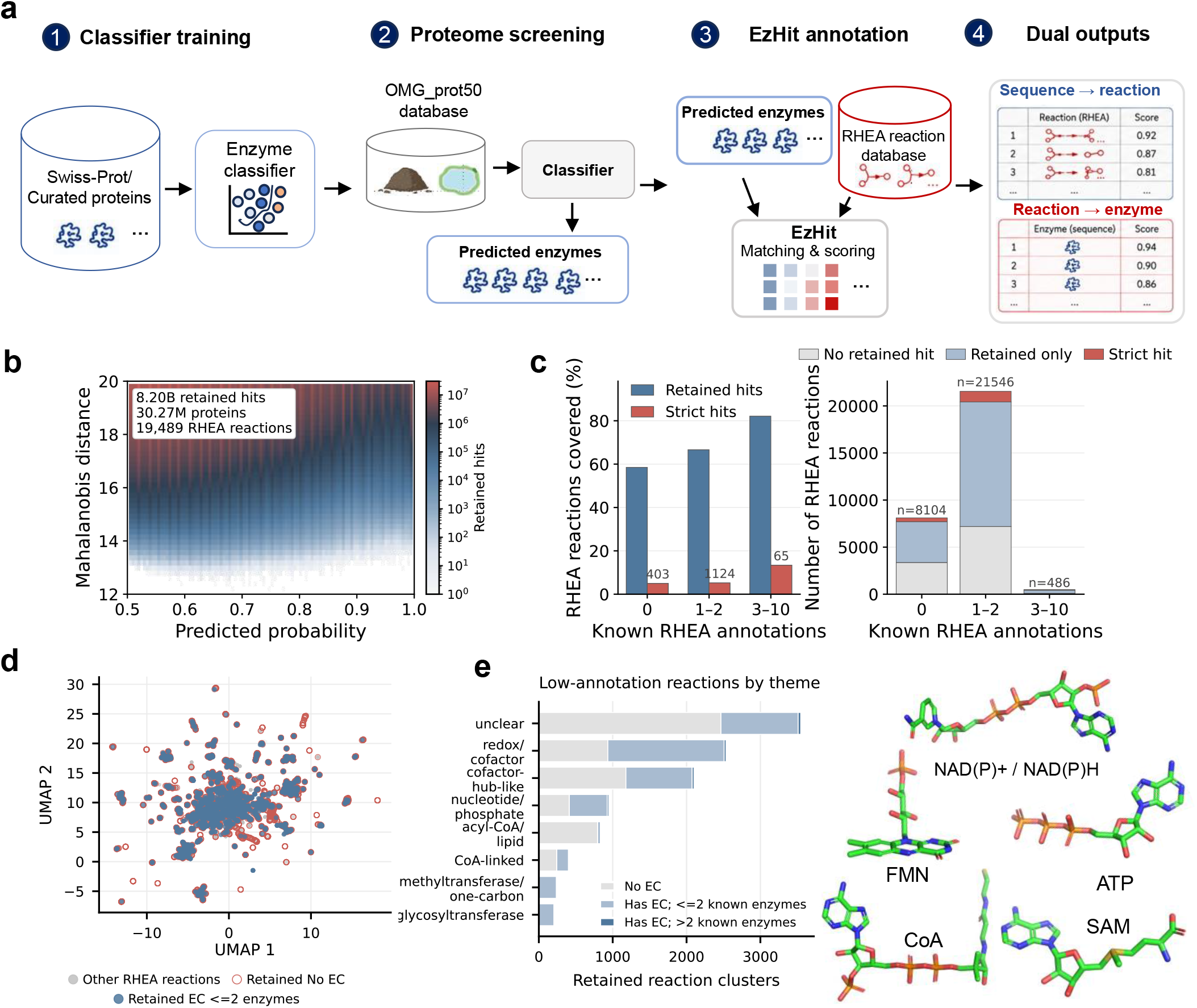
Metagenome scale EzHit screening maps candidate enzymes across underannotated RHEA chemistry. **a**, Workflow for large-scale screening. A sequence-level enzyme classifier trained on curated Swiss-Prot proteins was applied to OMG_Prot50, and predicted enzyme-like sequences were subsequently screened against RHEA using EzHit. **b**, Confidence landscape of retained enzyme–reaction associations across predicted probability and Mahalanobis distance. Retained hits were defined by predicted probability > 0.5 and Mahalanobis distance < 20. **c**, Coverage of RHEA reactions grouped by the number of existing linked enzyme annotations. Retained and strict candidates are shown, with strict hits defined by predicted probability ≥ 0.8 and Mahalanobis distance ≤ 16. **d**, UMAP projection of RHEA reaction representations. Grey points denote other RHEA reactions; red points denote retained reactions without EC annotation and blue points denote retained EC-annotated reactions with no more than two known enzymes. **e**, Distribution of retained reaction clusters across broad biochemical themes and existing annotation levels. Representative cofactors associated with major reaction themes are shown.

Low-annotation reactions supported by EzHit were distributed throughout the broader RHEA representation space (**Fig. 5d**). Their biochemical composition was also diverse. Reaction-theme analysis identified substantial representation of redox and cofactor-dependent transformations, nucleotide- and phosphate-associated chemistry, acyl-CoA and lipid-related reactions, and one-carbon transfer chemistry (**Fig. 5e**). Together, these analyses show that sparsely annotated reactions supported by EzHit span a diverse range of biochemical classes.

### EzHit extends enzyme retrieval beyond close se uence homology

The large-scale screen also revealed high-confidence candidates that extended beyond close sequence homologues of characterized enzymes. Across the metagenomic candidate set, EzHit confidence remained high for a substantial number of proteins with limited similarity to their closest Swiss-Prot matches. In particular, 209 candidates had less than 30% sequence identity while remaining above the high-confidence threshold (**Fig. 6a**). These results show that EzHit can prioritize high-confidence enzyme–reaction associations beyond close sequence homology. We further examined whether candidates with limited sequence similarity retained structural similarity to known enzymes. Comparison of predicted candidate structures with experimentally determined PDB structures showed that many low-identity candidates matched established enzyme folds with high TM-scores (**Fig. 6b**). Representative superpositions illustrate several cases in which the overall enzyme architecture was well conserved despite substantial sequence divergence (**Fig. 6c**). The sequence and structural analyses therefore identify a group of metagenomic proteins that are distinct from close homologues but remain consistent with known enzyme architectures.

**Figure 6.**
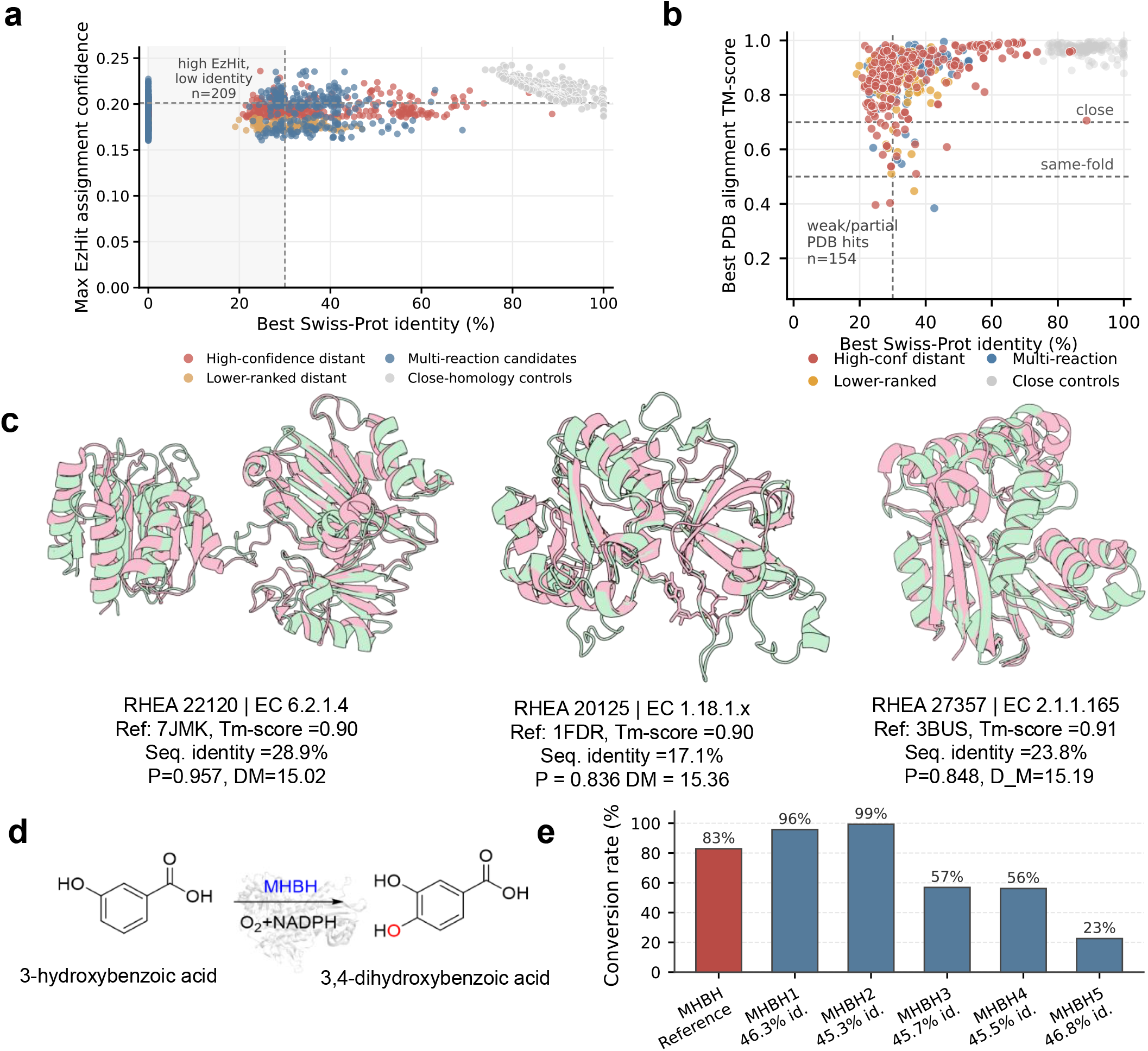
EzHit prioritizes se uence divergent candidates and identifies active enzymes for a sparsely annotated reaction. **a**, Relationship between EzHit assignment confidence and sequence similarity to characterized Swiss-Prot proteins for candidates from the metagenomic screen. The dashed vertical line marks 30% sequence identity and the dashed horizontal line marks the confidence threshold. A subset of 209 candidates retained high EzHit confidence despite low sequence identity to characterized proteins. **b**, Structural assessment of folded candidates against experimentally determined PDB structures. Each point is positioned by its best Swiss-Prot sequence identity and best PDB alignment TM-score. Dashed horizontal lines mark approximate same-fold and close-structure regimes. A subset of candidates with limited sequence identity retained strong structural similarity to known enzyme folds. **c**, Representative structural comparisons of selected candidates and their closest same-reaction or same-EC reference structures. Candidate structures are shown in pale green and reference structures in pink. Labels report the corresponding RHEA reaction, EC annotation, reference PDB structure, sequence identity, EzHit predicted probability and Mahalanobis distance. **d**, Reaction scheme for experimental validation of RHEA:11480, an NADPH-dependent hydroxylation converting 3-hydroxybenzoate to 3,4-dihydroxybenzoate. MHBH is the characterized reference enzyme currently linked to this reaction. **e**, Experimental conversion of 3-hydroxybenzoate by MHBH and five EzHit-prioritized metagenomic candidates. Bars show substrate conversion from a single measurement for each enzyme. Sequence identities shown below MHBH1–MHBH5 are calculated relative to MHBH. All five predicted candidates showed measurable conversion, ranging from 22.6% to 99.3%.

To experimentally test whether the large-scale screen could recover active enzymes, we selected RHEA:11480, an NADPH-dependent hydroxylation converting 3-hydroxybenzoate to 3,4-dihydroxybenzoate, or protocatechuate (Fig. 6d). Protocatechuate is a useful aromatic intermediate and a precursor for value-added compounds including vanillic acid and vanillin.^31–32^ At the time of analysis, RHEA:11480 was linked to only one UniProtKB protein, m-hydroxybenzoate hydroxylase (MHBH), making it a suitable example of a sparsely annotated activity. Five metagenomic candidates were selected using EzHit score, geometric confidence and supporting sequence and structural evidence. The candidates shared 45.3–46.8% sequence identity with MHBH, representing moderate sequence divergence from the reference enzyme.

All five candidates showed measurable substrate conversion in the target reaction (Fig. 6d). MHBH1 and MHBH2 reached 95.8% and 99.3% conversion, respectively, compared with 82.9% for the characterized MHBH reference, while MHBH3, MHBH4 and MHBH5 showed conversions of 56.9%, 56.2% and 22.6%, respectively. Although these measurements do not by themselves establish differences in catalytic efficiency, they demonstrate that EzHit-guided screening can recover active metagenomic enzymes for a sparsely annotated reaction. Together with the sequence and structural analyses, this result shows how atlas-scale retrieval can narrow a large metagenomic sequence space to experimentally testable enzyme candidates.

## Discussion

The performance of EzHit raises a broader question about how much structural information is needed for large-scale enzyme retrieval. Enzyme catalysis ultimately depends on three-dimensional structure, but predicted structures can contain local errors and may not fully represent ligand-dependent conformations or interactions.^33^ We also found that increasing dropout improved both performance and training stability. This may reflect the tendency of enzyme–reaction models to learn dataset-specific patterns from uneven and incomplete biochemical annotations, which remain a known limitation of enzyme databases.^34–35^ The results therefore suggest that effective regularization may be as important as increasing model complexity when learning enzyme–reaction relationships from current datasets.

Our uncertainty analysis highlights a high prediction score does not necessarily imply high reliability when a query differs from the training distribution, a general problem in machine learning under distribution shift.^36^ In our experiments, probability-based uncertainty measures and ensemble disagreement did not consistently identify such cases, and evidential learning reduced retrieval performance. In contrast, Mahalanobis distance in the learned feature space provided a more useful confidence signal. Feature-space distance has previously been used to identify out-of-distribution samples [8], and our results extend this idea to enzyme–reaction retrieval. They suggest that the position of a new enzyme–reaction pair relative to the training distribution can provide useful information beyond its prediction score alone. This distinction is particularly important in enzyme mining, where only a small fraction of computational candidates can usually be tested experimentally.

The OMG_Prot50 × RHEA screen showed how these features can be used in a practical discovery setting. EzHit identified candidate sequences for hundreds of reactions with few or no linked enzyme annotations and showed that these reactions span diverse biochemical classes. The candidate space could then be narrowed using prediction score, confidence, sequence similarity and structural evidence. This analysis identified candidates beyond the closest sequence matches to known enzymes, while many retained structural similarities to established enzyme folds. Experimental testing of selected candidates for a sparsely annotated hydroxylation further supported the catalytic activity predicted by the screen. The value of the billions of enzyme–reaction associations generated by EzHit therefore lies not in the predictions themselves, but in reducing a very large sequence space to a small set of experimentally testable candidates and expanding the enzyme sequence space associated with poorly characterized reactions.

The current study also suggests several directions for future development. The strong performance of EzHit shows that sequence–reaction representations alone can support effective large-scale enzyme retrieval and, in the benchmark, settings examined here, can outperform approaches that additionally incorporate protein structural information. This observation suggests that the key challenge may not simply be whether structural information is included, but how sequence, structure and reaction information are integrated. Future work could therefore explore more effective multimodal representations that capture complementary structural signals without weakening the information already encoded in protein sequence representations or compromising large-scale screening efficiency. Confidence estimation could also be further developed for increasingly diverse sequence and reaction spaces, while iterative experimental validation may help refine retrieval models and expand enzyme annotations for poorly characterized reactions.

## Conclusions

In this work, we developed EzHit as a scalable framework for bidirectional enzyme–reaction retrieval and applied it from benchmark evaluation to metagenomic-scale biocatalyst discovery. EzHit achieved strong retrieval performance without requiring explicit protein structural inputs or enzyme–substrate complex modelling, while feature-space confidence provided additional information for prioritizing predictions beyond the training distribution. Its computational efficiency enabled screening of nearly one trillion enzyme–reaction pairs across OMG_Prot50 and RHEA and generated ranked candidates for many sparsely annotated reactions. Experimental testing of five candidates for the sparsely annotated NADPH-dependent hydroxylation of 3-hydroxybenzoate to protocatechuate showed measurable activity in all cases. These results demonstrate that large-scale enzyme–reaction retrieval can connect expanding protein sequence collections with experimentally testable biocatalyst candidates and support reaction-guided enzyme discovery.

## Methods

### Dataset

To train and evaluate the proposed model, we utilized the datasets originally curated in the EnzymeCAGE framework.^12^ The datasets were primarily sourced from the Rhea database, employing a time-based data partitioning strategy to ensure rigorous evaluation of the model’s generalization capabilities.

### Training Dataset

The training data was derived from the Rhea database^6^ snapshot before uly 12, 2023 (version 128). Entries lacking UniProt identification numbers or sequence information were removed. Additionally, reactions involving substrates with more than 150 atoms were excluded, redundant molecules present in both reactants and products were eliminated, and carbon-containing substrates were standardized from ionic to charge-balanced molecular forms. Proteins with sequence lengths exceeding 1000 were also filtered out for computational efficiency. Because the Rhea database contains only positive enzyme-reaction pairs, negative samples were generated at an approximate positive-to-negative ratio of 1:5. Negative enzymes for a target reaction were selected by searching for similar reactions in the database and extracting their associated enzymes, ensuring true positive enzymes were excluded. The final curated training dataset includes 1,513,592 enzyme-reaction pairs.

### Internal Test Sets

To benchmark the model against unseen enzymes and reactions, two internal test sets were constructed using data incorporated into Rhea between uly 12, 2023, and February 5, 2025. To prevent data leakage, any reactions exhibiting a similarity score greater than 0.9 to reactions in the training set were excluded.

#### Enzyme 405 Dataset

This set was designed to evaluate function prediction for unseen enzymes. It comprises 295 reactions and 405 enzymes whose sequences were absent from the training set. Negative enzymes were constructed by identifying the ten most similar reactions from the uly 2023 Rhea snapshot and assigning their corresponding enzymes as negatives. This resulted in a total of 15,921 enzyme-reaction pairs (961 positives and 14,960 negatives).

#### Orphan 335 Dataset

This set evaluates the model’s capability in reaction de-orphaning. It contains 335 orphan reactions that had no annotated enzymes in the uly 2023 training snapshot but were subsequently annotated by February 2025.

### External Domain Specific Test Sets

To assess predictive performance within specific, highly diverse enzyme families, three external test sets were utilized: cytochrome P450s,^22^ terpene synthases,^23^ and phosphatases.^24^ For all three datasets, reactions with a similarity score greater than 0.9 to those in the training set were removed to prevent data leakage. Specifically, the cytochrome P450s dataset was formulated using a many-to-many data construction approach containing 191 reactions and 490 enzymes, yielding 93,590 candidate enzyme-reaction pairs of which 318 are true positives. Similarly, the terpene synthases dataset was constructed using the same many-to-many approach, including 92 reactions and 1,093 enzymes, resulting in 100,556 enzyme-reaction pairs. In contrast to the other families, the phosphatases dataset inherently contained experimentally validated negative samples, thereby negating the need for manual negative sample generation. This final dataset comprises 95 reactions and 178 enzymes, totaling 16,910 pairs.

### Model Architecture

To accurately and efficiently predict the catalytic specificity between an enzyme and a chemical reaction, we developed a Late-Fusion Cross-Encoder architecture named HybridPairKAN.^21^ Unlike computationally prohibitive early-fusion models that require recalculating sequence-reaction embeddings for every possible pair, our architecture relies on pre-computed, modality-specific high-dimensional representations. These features are then fused and mapped to a catalytic probability score through a hybrid network combining a Multilayer Perceptron (MLP) bottleneck and a Kolmogorov-Arnold Network (KAN).

### Input Representations

The model takes three distinct high-dimensional embeddings as input. Specifically, the enzyme sequence embedding (*E*) is extracted using the pre-trained ProtT5 language model,^18^ which captures the global biochemical and structural properties of the enzyme (*D*_*T5*_ = 1024). For the chemical reaction, we utilize a Differential Reaction Fingerprint (*D*_*DRFP*_) to encode the overall reaction dynamics and atomic environment changes between reactants and products (*D*_*DRFP*_ = 2048),^19^ alongside a Reactant Fingerprint (*R*) based on the Extended-Connectivity Fingerprint (ECFP/Morgan Fingerprint) to exclusively encode the substrate topology (*D*_*ECFP*_ = 2048).^20^ To mitigate the scale discrepancies among these features derived from different pre-trained encoders, we apply independent Layer Normalization to each modality before fusion:

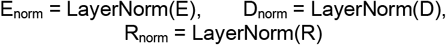

The normalized vectors are then concatenated into a single fused feature vector 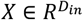, where *D*_*in*_ = *D*_*T5*_ *D*_*DRFP*_ *D*_*ECFP*_.

### Feature Compression and Bottleneck Layer

Directly feeding the high-dimensional concatenated vector into a complex non-linear network can lead to optimization instabilities and exorbitant memory costs. To address this, we introduce an MLP-based compressor module. This bottleneck layer projects the concatenated features into a lower-dimensional, dense semantic space:

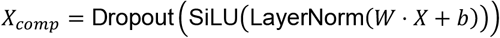

where 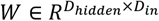 is a learnable weight matrix, *D*_*hidden*_ represents the bottleneck dimension, and SiLU is the Sigmoid Linear Unit activation function. This module efficiently filters out redundant noise while capturing the initial cross-modal interactions.

### Kolmogorov Arnold Network (KAN) for Catalytic Scoring

For the final predictive reasoning, we replace traditional dense layers (MLPs) with a Kolmogorov-Arnold Network (KAN). Unlike standard MLPs that use fixed activation functions on nodes,^21^ KANs parameterize learnable 1D non-linear activation functions on the edges (weights) of the network. This fundamental shift allows the model to learn highly complex and potentially interpretable non-linear relationships between the compressed enzyme and reaction features. In our architecture, the KAN maps the compressed feature vector *X*_*comp*_ to a scalar catalytic logit. For a given input scalar *x* on a specific edge, the 1D activation function *ϕ*(*x*) is formulated as the combination of a base residual transformation and a learnable B-spline curve:

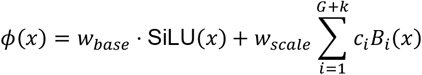

where SiLU(x) serves as the base activation function with a learnable weight *w*_*base*_. The non-linear spline component is constructed using B-spline basis functions *B*_*i*_(*x*) of order *k* (set to 3, cubic splines), defined over a grid of size *G* (set to 5). The parameters c_*i*_ represent the learnable spline coefficients, and *w*_*scale*_ is an independent scaling factor (spline_scaler) to stabilize training.

### Dynamic Grid Adaptation

A critical challenge in spline-based networks is ensuring that the bounded B-spline grids effectively cover the active range of the dynamic input features. To address this, our KAN implementation employs a dynamic grid updating mechanism (update_grid). During the forward pass, the knot positions of the B-splines are periodically adjusted to fit the empirical distribution of the incoming activation values. The updated grid is calculated as a weighted mixture of a uniform spatial grid and an adaptive, density-based grid (computed via data quantiles), ensuring robust interpolation across dense feature regions while preventing boundary overflow.

### Optimization Objective and Sparsity Regularization

The model is trained end-to-end using a weighted Binary Cross-Entropy (BCE) loss combined with a KAN-specific sparsity regularization term. To counteract the extreme class imbalance typical of metagenomic datasets (where negative non-enzyme pairs vastly outnumber positive hits), we dynamically compute a positive weight (*α*_*pos*_ = *N*_*neg*_/*max*(1, *N*_*pos*_)) for the BCE loss.^37^

Furthermore, to encourage network sparsity and improve the potential interpretability of the learned spline functions, we introduce a regularization loss 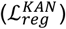 applied to the KAN layers. This term is an efficient approximation of the *L*_1_ norm of the spline activations, augmented with an entropy penalty:

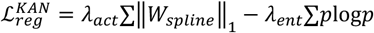

where ‖*W*_*spline*_‖_1_ represents the mean absolute values of the spline coefficients, and *p* denotes the normalized magnitude distribution of these weights. The total loss function optimized during training is defined as:

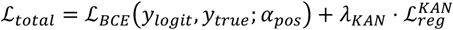

where *λ*_*KAN*_ controls the strength of the KAN regularization, effectively pruning redundant spline edges and mitigating overfitting. The final catalytic probability is obtained via a Sigmoid activation: *P* = *σ*(*y*_*logit*_) ∈ [0,1].

### Uncertainty Quantification (Q) Strategies

To systematically evaluate the reliability of our enzyme-reaction pair predictions and the model’s robustness against out-of-distribution (OOD) data, we implemented and compared multiple uncertainty quantification (U) strategies. These methods can be broadly categorized into traditional probability-based metrics and advanced feature-space geometric metrics. For consistency, all scores were converted so that larger values indicate higher uncertainty, unless otherwise specified.

### Probability based and Ensemble Strategies

*<*H3>Predictive Confidence

The standard baseline for uncertainty is derived directly from the maximum softmax probability (or sigmoid output) of the model.^26–27^ The uncertainty score is defined as:

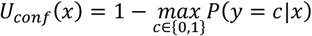

#### Ensemble Mutual Information (MI) & Predictive Entropy

To capture epistemic uncertainty, we utilized an ensemble strategy comprising *M* independent models.^25^ For a given input *x*, let 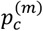 be the predicted probability for class *c* from the *m*-th model, and 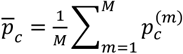 be the mean prediction. The Predictive Entropy evaluates the total uncertainty:

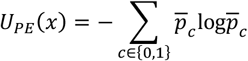

The Mutual Information (MI) captures the disagreement among the ensemble members, representing the epistemic uncertainty:

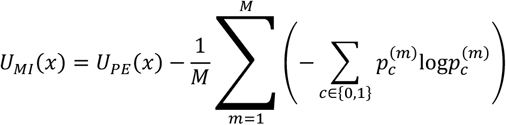

### Feature Space Geometric Strategies

Due to the highly non-linear nature of the Kolmogorov-Arnold Networks (KAN) utilized in our architecture, we hypothesized that OOD samples might suffer from “origin collapse” or manifold distortion, rendering traditional probability scores overconfident. Therefore, we extracted the penultimate bottleneck representations *h*(*x*) to compute distance-aware U metrics.

#### Negative Mahalanobis Distance

We fit a class-conditional Gaussian distribution to the training features by calculating the mean vector *μ*_*c*_ for each class *c* and a tied empirical covariance matrix Σ. The Mahalanobis distance for a test sample is calculated as:

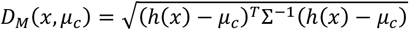

Counter-intuitively, our empirical analysis revealed an “inside-out” topology in the latent space, where highly confident and accurate predictions are pushed to the decision boundaries (larger distances), while ambiguous or OOD samples collapse toward the distribution mean.^28^ Thus, we utilized the negative distance to the closest class center as the uncertainty score:

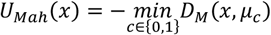

*Feature Activation Magnitude*. To further validate the “origin collapse” phenomenon, we assessed the magnitude of the activation vectors. We defined the uncertainty based on the negative *L*_2_ norm of the latent representation:

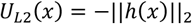

A higher score (lower absolute magnitude) suggests that the neural network lacks sufficient activation patterns recognizing the input, effectively flagging structurally dissimilar OOD entities.

#### K-Nearest Neighbors (KNN) Distance

To circumvent the assumption of global Gaussian distributions, we also implemented a non-parametric local density estimation. For a given test representation *h*(*x*), we calculate its Euclidean distance to the *K* nearest neighbors in the training set features.^38^ The uncertainty is the average distance:

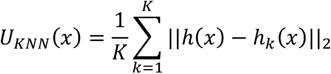

Where *h*_*k*_(*x*) denotes the *k*-th closest training sample. This metric accurately identifies isolated samples in the feature space as highly uncertain.

### Evaluation metrics and reaction similarity analysis

Model performance was evaluated using Top-*k* Success Rate, Enrichment Factor, Top-10 DCG, and ROC-AUC, depending on the corresponding benchmark setting. Top-*k* Success Rate measures whether at least one true positive candidate is retrieved within the top-*k* predictions, while Enrichment Factor and DCG evaluate the enrichment and ranking position of true positives among top-ranked candidates. ROC-AUC was used for binary classification settings. Reaction similarity was calculated following the Selezyme strategy using Morgan fingerprints and Tanimoto coefficients, considering both direct reactant-product matching and cross-matching. Detailed definitions are provided in **Supplementary Note 7**.

### Protein expression and purification

Target genes were cloned into the pET-28a expression vector and transformed into *Escherichia coli* BL21(DE3) competent cells. Single colonies were selected and cultured in LB medium. Seed cultures were inoculated at 10% (v/v) into fresh LB medium and grown at 37 °C with shaking at 200 rpm. When the culture reached an OD_600_ of 0.6–0.8, protein expression was induced by the addition of IPTG to a final concentration of 0.5 mM. The cultures were then incubated at 25 °C and 200 rpm for 20 h. Cells were harvested by centrifugation and used for subsequent protein purification.

His-tagged proteins were purified by immobilized metal affinity chromatography. Buffer A contained 20 mM imidazole, 500 mM NaCl and 20 mM Na_3_PO_4_, and buffer B contained 300 mM imidazole, 500 mM NaCl and 20 mM Na_3_PO_4_. Purified proteins were exchanged into 50 mM Tris–HCl buffer (pH 7.5) and stored in the same buffer before use in enzyme assays.

### In vitro activity assay

Enzymatic reactions were performed in a total volume of 400 μL containing 0.5 mM 3-hydroxybenzoic acid (3-HBA), 0.5 mM NADPH and 0.4 mg mL^-1^ purified enzyme in 50 mM Tris–HCl buffer (pH 7.5). Reactions were incubated at 30 °C with shaking at 900 rpm for 30 h. After incubation, reactions were quenched by adding an equal volume of methanol and were then subjected to HPLC analysis. For experimental validation of RHEA:11480, MHBH was used as the characterized reference enzyme, and five EzHit-prioritized metagenomic candidates were assayed under the same conditions.

### HPLC analysis and quantification

Reaction products were analyzed by high-performance liquid chromatography using a C18 column (5 μm, 4.6 × 250 mm). The flow rate was 1.0 mL min^-1^, the column temperature was maintained at 30 °C, the injection volume was 10 μL and the detection wavelength was 260 nm. Mobile phase A was acetonitrile and mobile phase B was ultrapure water containing 0.1% (v/v) trifluoroacetic acid.

The elution program consisted of 2% A and 98% B from 0 to 2.0 min, a linear increase to 40% A and 60% B from 2.0 to 15.0 min, a return to 2% A and 98% B at 15.1 min, and equilibration at 2% A and 98% B until 20.0 min.

Substrate conversion and product formation were quantified using external standard curves of 3-HBA and protocatechuic acid (PCA). Standard solutions ranging from 0.015625 to 2 mM were analyzed by HPLC under the same conditions. Linear calibration curves were generated from peak area versus concentration and were used to calculate substrate conversion and product yield in enzymatic reactions.

## DATA AVAILABILITY

All processed datasets generated or analyzed in this study are available at [[LINK](https://doi.org/10.5281/zenodo.20278477)]. Raw reaction annotations were obtained from RHEA at [[LINK](https://www.rhea-db.org/help/download)], and protein sequences were obtained from [[LINK](https://huggingface.co/datasets/tattabio/OMG_prot50)]. All remaining data supporting the conclusions of this work are included in the main text and supplementary materials. The large-scale OMG_Prot50 × RHEA screening results generated in this study are available at [[LINK](https://huggingface.co/datasets/deanluo/EzHit-large-scale-screening)].The EzHit source code is available at [[LINK](https://github.com/ld139/EzHit)]. The repository includes scripts for preprocessing enzyme–reaction datasets, training and fine-tuning EzHit models, performing enzyme retrieval and annotation, calculating uncertainty scores, and reproducing the analyses and figures reported in this study. Pretrained checkpoints and example files are available at [[LINK](https://huggingface.co/deanluo/EzHit)]. A user-friendly Google Colab workflow for prediction, retrieval, and optional fine-tuning is provided at [[LINK](https://colab.research.google.com/github/ld139/EzHit/blob/main/colab/EZHit_FineTune_Colab.ipynb)]. The code is released under the MIT license.

## ASSOCIATED CONTENT

### Supporting Information

Details of model architectures, additional experimental results, training parameters, enzyme sequences (PDF)

## ACKNOWLEDGMENTS

This work has been supported by Scientific Research Innovation Capability Support Project for Young Faculty (ZYGXON SKYCXNLZCXM-B6), National Natural Science Foundation of China (22121001), and Fundamental Research Funds for the Central Universities (20720240124).

