## Supplementary material for "Scalable Enzyme–Reaction Retrieval Enables Metagenomic Biocatalyst Discovery": SI

### **Supplementary Notes**

#### **Supplementary Note 1 | Baseline models**

To provide a comprehensive evaluation of EzHit, we compared it with representative baseline models for enzyme–reaction or enzyme–substrate association prediction, including ESP^1^, CLIPZyme^2^, and EnzymeCAGE ^3^. These models were selected because they represent different modeling strategies for connecting enzyme sequences with chemical substrates or reactions, ranging from sequence–molecule matching to contrastive representation learning and enzyme–reaction retrieval. All baseline models were evaluated under the same retrieval setting as EzHit using identical query reactions, candidate enzyme pools, and evaluation metrics.

ESP^1^ is a sequence- and substrate-based enzyme–substrate prediction model. It uses enzyme sequence information together with molecular representations of candidate substrates to estimate whether an enzyme can act on a given substrate. In our benchmark, ESP was used as a representative substrate-level prediction baseline for evaluating whether sequence–molecule compatibility alone is sufficient for orphan-reaction retrieval.

CLIPZyme^2^ is a contrastive representation learning model designed to align enzyme and chemical reaction information in a shared embedding space. By learning matched enzyme–reaction pairs, CLIPZyme enables retrieval-style prediction through similarity scoring between enzyme and reaction embeddings. We included CLIPZyme as a baseline to assess the performance of contrastive enzyme–reaction representation learning in sparse and orphan-reaction settings.

EnzymeCAGE^3^ is a recent enzyme–reaction retrieval model that integrates enzyme sequence representations with reaction-level information to identify candidate enzymes for biochemical reactions. Because EnzymeCAGE is specifically designed for enzyme–reaction matching and retrieval, it was used as the primary state-of-the-art baseline in our comparative analyses. In addition to the full EnzymeCAGE model, we also compared EzHit against EnzymeCAGE in fine-tuning experiments to evaluate whether EzHit maintains an advantage under enzyme-family-specific adaptation.

For all baseline models, we followed the recommended inference or training protocols from the original implementations whenever available. For ClipZyme, because retraining is computationally expensive and the released model weights are large, we directly used the provided pretrained checkpoint for inference. Reaction entries that could not be processed by the released ClipZyme pipeline were excluded from the corresponding evaluation. For ESP, we retrained the model on the same training splits using the default hyperparameters, and reported the results from five independent random seeds. For EnzymeCAGE, we used the five pretrained checkpoints released by the original authors and reported the averaged performance across these seeds. We did not include EZSpecificity^4^ in the benchmark comparison because it requires docked enzyme–substrate poses as input. As docking pose generation can introduce additional uncertainty unrelated to the predictive model itself, including this model would make it difficult to separate model performance from pose-generation quality. To ensure a fair comparison, all applicable baselines were evaluated using the same candidate pools as EzHit whenever possible. Performance was assessed using Top-k Success Rate, Enrichment Factor, Top-10 DCG, and ROC-AUC, depending on the corresponding benchmark setting.

#### **Supplementary Note 2 | Theoretical formulation and implementation of evidential deep learning**

To evaluate uncertainty quantification strategies without relying on model ensembling, we implemented Evidential Deep Learning (EDL) as an additional uncertainty baseline.^5^ Unlike standard classifiers that directly output deterministic class probabilities, EDL treats the network outputs as non-negative evidence values that parameterize a Dirichlet distribution over class probabilities. This enables the model to estimate predictive probability and epistemic uncertainty within a single forward pass.

In our implementation, the evidential model was built on the same enzyme–reaction feature representation used by the KAN-based classifier. Specifically, each enzyme–reaction pair was represented by three feature types: enzyme sequence embeddings, DRFP reaction fingerprints,^6^ and reactant Morgan fingerprints.^7^ These features were individually normalized, concatenated, and projected into a shared hidden representation before being passed to a KAN module. ^8^ The final KAN output dimension was set to two, corresponding to the evidence values for the negative and positive classes. A Softplus activation was applied to ensure non-negative evidence:

$$e=\left[ e_{0},e_{1} \right]$$

The evidence values were converted into Dirichlet concentration parameters:

$$\alpha_{k}=e_{k}+1$$

and the total evidence was defined as:

$$S=\sum\alpha_{k}$$

where *k* =2 for binary classification. The predicted probability for class $k$was computed as the expectation of the Dirichlet distribution:

$$p_{k}=\frac{\alpha_{k}}{S}$$

For enzyme–reaction pair scoring, the positive-class probability *p*_1_ was used as the prediction score. The epistemic uncertainty was defined as:

$$u=\frac{K}{S}$$

Therefore, samples with lower total evidence have higher epistemic uncertainty, whereas samples with stronger accumulated evidence have lower uncertainty.

The model was trained using an evidential objective consisting of an expected mean squared error term and a KL-divergence regularization term. For a one-hot encoded label $y$, the expected MSE term was:

$$\mathcal{L}_{\mathrm{MSE}}=\sum_{k=1}^{K} \left[ \left( y_{k}-\frac{\alpha_{k}}{S} \right)^{2}+\frac{\alpha_{k}(S-\alpha_{k})}{S^{2}(S+1)} \right]$$

The KL term regularized evidence assigned to incorrect classes by comparing the adjusted Dirichlet distribution with a uniform Dirichlet prior:

$$\mathcal{L}_{\mathrm{EDL}}=\mathcal{L}_{\mathrm{MSE}}+\lambda_{\mathrm{EDL}}\mathrm{KL}\left[ \mathrm{Dir}(\tilde{\alpha})\parallel\mathrm{Dir}(1) \right]$$

where $\tilde{\alpha}=y+(1-y)\alpha$, and $\lambda_{\mathrm{EDL}}$was set to 0.1 in our implementation. In addition, the KAN regularization loss was included during training to maintain consistency with the standard KAN training protocol. The final training objective was:

$$\mathcal{L=}\mathcal{L}_{\mathrm{EDL}}+\lambda_{\mathrm{KAN}}\mathcal{L}_{KAN-reg}$$

with $\lambda_{\mathrm{KAN}}=1\times{10}^{-4}$. During evaluation, EDL uncertainty was assessed by retaining samples with epistemic uncertainty below a predefined threshold and recalculating ranking and enrichment metrics on the retained subset. The EDL-KAN baseline used the same input feature types, reaction preprocessing procedure, and evaluation protocol as the standard EzHit KAN model, while several training hyperparameters were adjusted for stable evidential optimization, as summarized in **Supplementary Table 6**.

#### **Supplementary Note 3 | Performance degradation and out-of-distribution failure of EDL**

Although EDL provides a principled framework for single-pass uncertainty estimation, our empirical results showed that replacing the standard KAN classifier with the evidential KAN led to a substantial reduction in baseline predictive performance on both the Enzyme-405 and OOD Orphan-335 benchmarks (**Supplementary Fig. 5**). This indicates that the evidential training objective introduced a non-negligible optimization trade-off in our enzyme–reaction ranking setting.

A possible explanation is that the EDL objective simultaneously encourages accurate classification and conservative evidence assignment. While this design is useful for uncertainty estimation, it can conflict with the representation learning requirement of the KAN model, especially for complex enzyme–reaction decision boundaries. The KL regularization term penalizes excessive evidence for difficult or incorrectly classified samples, which may restrict the model from forming highly discriminative decision boundaries required for accurate candidate ranking.

We further evaluated whether EDL-derived epistemic uncertainty could improve prediction reliability by filtering out samples with high uncertainty. In this setting, the positive-class probability $p_{1}$was used as the ranking score, while $u=2/S$was used as the epistemic uncertainty score. Samples with uncertainty below the predefined threshold were retained for the EDL-filtered evaluation. However, the resulting confidence-based retention curves did not provide robust improvement under OOD conditions (**Supplementary Fig. 5**), suggesting that output-level evidence was insufficient to consistently identify structurally novel enzyme–reaction pairs.

These observations motivated the use of post-hoc geometric uncertainty quantification based on latent-space Mahalanobis distance. Unlike EDL, this strategy does not modify the training objective of the base KAN model and therefore avoids the observed performance trade-off. At the same time, latent-space distance directly measures how far a test enzyme–reaction pair is from the training distribution, making it more suitable for detecting OOD enzyme–reaction candidates.

#### **Supplementary Note 4 | Massive-scale bipartite inference and uncertainty-based filtering**

Large-scale screening of OMG_Prot50-derived enzyme candidates against the RHEA reaction library requires evaluating an extensive enzyme–reaction Cartesian product. Directly materializing this prediction matrix is computationally infeasible because of GPU memory limits and disk I/O overhead. To address this challenge, we implemented a batched bipartite inference pipeline that combines GPU-accelerated tensor operations, early uncertainty-aware filtering, and streaming output.

For each candidate enzyme shard, protein embeddings were loaded from the corresponding cache, while RHEA reaction features were precomputed using DRFP reaction fingerprints and reactant Morgan fingerprints. The RHEA feature tensors were kept on GPU memory during inference to avoid repeated feature construction. The trained KAN-based EzHit model was then used to score enzyme–reaction pairs in a block-wise manner. Specifically, each protein batch was paired with a reaction batch through tensor-level expansion, allowing the model to evaluate a local Cartesian product without constructing the full enzyme–reaction matrix.

In parallel with prediction, we computed a latent-space Mahalanobis distance for each enzyme–reaction pair. The positive-class mean vector and inverse covariance matrix estimated from the training distribution were loaded before inference. For each pair, the model generated both a prediction logit and a latent representation. The predicted probability was obtained by applying a sigmoid function to the logit, while the Mahalanobis distance was calculated from the latent representation relative to the positive-class training distribution.

To control memory usage, filtering was applied immediately after each inference block. Only enzyme–reaction pairs satisfying both criteria, predicted probability greater than 0.5 and Mahalanobis distance lower than 20.0, were retained. For each retained pair, only the protein identifier, RHEA identifier, predicted probability, and Mahalanobis distance were written to disk. The retained results were streamed directly into compressed Parquet files, which avoided accumulating the full prediction matrix in memory.

This pipeline enabled large-scale screening over billions of potential enzyme–reaction pairs while maintaining a bounded memory footprint. The resulting high-confidence, in-distribution candidate pairs were used for downstream reaction-level prioritization and candidate selection. Importantly, these retained enzyme–reaction associations should be interpreted as computationally prioritized hypotheses rather than experimentally confirmed functional annotations.

#### **Supplementary Note 5 | Enzyme candidate pre-filtering for OMG_Prot50-scale screening**

To reduce the computational cost of large-scale enzyme–reaction screening, we introduced a sequence-level enzyme/non-enzyme pre-filtering step before applying EzHit to the OMG_Prot50 database. This classifier was used only to remove unlikely non-enzyme sequences and was not used as the final enzyme–reaction association predictor.

The enzyme/non-enzyme classifier was trained using Swiss-Prot proteins. Proteins with annotated EC numbers were labeled as enzymes, whereas proteins without EC annotations were labeled as non-enzymes. Each protein sequence was represented by a 1024-dimensional ProtT5-derived sequence embedding. To preserve the original enzyme/non-enzyme class distribution during model development, the dataset was split into training and validation sets using a stratified 80/20 split based on the binary enzyme label. The validation set was used to monitor convergence and select the final checkpoint, rather than to serve as an independent downstream benchmark.

The classifier used a dual-head neural network architecture consisting of a shared encoder, a projection head, and a binary classification head. The projection head was optimized with supervised contrastive loss to improve enzyme/non-enzyme separation in the latent space, while the classification head was optimized with weighted binary cross-entropy loss to predict enzyme probability. The final objective combined these two terms:

$$L=0.3L_{supcon}+0.7L_{BCE}$$

Class imbalance was handled by assigning a positive-class weight according to the ratio of non-enzyme to enzyme samples. The model was trained with early stopping based on validation loss. Training converged within 14 epochs, at which point early stopping was triggered after five consecutive epochs without further validation-loss improvement. At the final epoch, the model achieved a validation accuracy of 0.9882, with a classification loss of 0.0200. The overall loss remained dominated by the supervised contrastive component because this term operates on pairwise feature separation and has a different numerical scale from the binary classification loss. The checkpoint with the lowest validation loss was retained as the final enzyme/non-enzyme classifier for downstream filtering. Detailed training hyperparameters are summarized in **Supplementary Table 5**.

Before enzyme probability prediction, the OMG_Prot50 database was globally cleaned and repartitioned. Sequences containing ambiguous or non-standard amino acid characters, including X, U, Z, O, and B, were removed, and only sequences with lengths between 100 and 1500 amino acids were retained. The remaining sequences were deduplicated at the sequence level while preserving one original identifier for each unique sequence. To balance the inference workload, the deduplicated sequences were sorted by sequence length and redistributed into 32 length-balanced Parquet shards using a round-robin strategy.

Each cleaned OMG_Prot50 shard was then processed independently. Protein sequences were converted into ProtT5-compatible spaced amino acid strings and encoded with the ProtT5 encoder. Sequence-level embeddings were obtained by attention-mask-based mean pooling over the final hidden states. To improve throughput, dynamic batching was performed according to a maximum token budget, and sequences were sorted by length within each batch to reduce padding overhead. The trained enzyme/non-enzyme classifier assigned an enzyme probability score to each sequence, and sequences with predicted probability greater than 0.5 were retained as putative enzyme candidates. The retained protein identifiers, sequences, enzyme probabilities, and ProtT5 embeddings were written to Parquet files using streaming output. The overall pre-filtering workflow is illustrated in **Supplementary Fig. 10**.

#### **Supplementary Note 6 | Inference runtime comparison between EzHit and EnzymeCAGE**

To evaluate the computational efficiency of EzHit relative to a structure-aware baseline, we benchmarked the inference runtime of EzHit and EnzymeCAGE under the Enzyme-405 test setting. All runtime measurements were performed on the same NVIDIA GeForce RTX 3090 GPU. Each model was first executed once as a warm-up run, followed by five repeated timed inference runs. CUDA synchronization was applied before and after each timed run. The detailed runtime results are summarized in **Supplementary Table 8**.

For EzHit, runtime was measured using the seed42 checkpoint and precomputed input features, including enzyme sequence embeddings, DRFP reaction fingerprints, and reactant Morgan fingerprints. Therefore, the reported EzHit runtime corresponds to cached-feature model inference from input tensors to prediction scores, excluding one-time feature generation and disk writing. For EnzymeCAGE, runtime was measured using its official DataLoader-based *model.evaluate* inference loop, which reflects the standard inference procedure of the released implementation with graph-structured inputs.

To avoid bias from different batch sizes, we first compared EzHit and EnzymeCAGE under the same batch size of 256. Under this controlled setting, EzHit processed 15,921 enzyme–reaction pairs in 0.101±0.003s, corresponding to 157,696 pairs/s and 6.34 s per million pairs. In comparison, EnzymeCAGE processed 15,611 pairs in 39.36±0.97s, corresponding to 397 pairs/s and 2521.00 s per million pairs. Thus, under the same batch size, EzHit achieved approximately 398-fold higher inference throughput than EnzymeCAGE.

We also evaluated EzHit under its practical high-throughput setting with a batch size of 8192, which is feasible because EzHit operates on dense cached tensor features rather than graph-structured inputs. In this setting, EzHit processed 15,921 pairs in 0.0749±0.0003s, corresponding to 212,631 pairs/s and 4.70 s per million pairs. Compared with the official EnzymeCAGE inference setting, this corresponds to an approximately 536-fold higher practical inference throughput.

The difference in the number of evaluated pairs arises because EnzymeCAGE requires graph-structured protein features and therefore excludes entries without the required geometric or node-level features, whereas EzHit operates on cached tabular feature tensors. We therefore report both the number of evaluated pairs and normalized throughput in **Supplementary Table 8**. Feature and dataset setup time was not included in the inference-only comparison. This benchmark should be interpreted as a comparison of practical inference throughput under each model’s input representation, with an additional same-batch-size control to ensure fair comparison.

#### **Supplementary Note 7 | Evaluation metrics and reaction similarity calculation**

**Evaluation metrics:**

For enzyme–reaction retrieval tasks, candidate enzymes or reactions were ranked according to the predicted EzHit score. Model performance was evaluated using Top-$k$ Success Rate, Enrichment Factor, and Top-10 Discounted Cumulative Gain, depending on the corresponding benchmark setting.

Top-$k$ Success Rate measures whether at least one true positive candidate is retrieved within the top $k$ranked predictions for each query:

$$\mathrm{Top}\text{-}k\mathrm{Success}=\frac{1}{N}\sum_{i=1}^{N} \mathbb{I}(r_{i}\leq k)$$

where $N$is the number of queries, $r_{i}$is the best rank of any true positive candidate for query $i$, and $\mathbb{I}(\cdot)$is the indicator function.

Enrichment Factor at the top $p\%$evaluates whether true positives are enriched among the highest-ranked candidates:

$$\mathrm{EF}_{p\%}=\frac{n_{\mathrm{TP},p\%}/n_{p\%}}{n_{\mathrm{TP},\mathrm{total}}/n_{\mathrm{total}}}$$

where $n_{\mathrm{TP},p\%}$is the number of true positives in the top $p\%$ranked candidates, $n_{p\%}$is the number of candidates in this top-ranked subset, $n_{\mathrm{TP},\mathrm{total}}$is the total number of true positives, and $n_{\mathrm{total}}$is the total number of candidates.

Top-10 Discounted Cumulative Gain was used to evaluate whether true positives were ranked near the top of the candidate list:

$$DCG@10=\sum_{j=1}^{10} \frac{\mathrm{rel}_{j}}{\log_{2}(j+1)}$$

where $\mathrm{rel}_{j}$is 1 if the candidate ranked at position $j$is a true positive and 0 otherwise.

For binary classification settings, receiver operating characteristic area under the curve was used:

$$\mathrm{ROC}\text{-}\mathrm{AUC}=\int_{0}^{1} \mathrm{TPR}(\mathrm{FPR})\text{ }d\mathrm{FPR}$$

ROC-AUC measures the threshold-independent ability of a model to assign higher scores to positive enzyme–reaction pairs than to negative pairs.

**Reaction similarity calculation**

Reaction similarity was calculated following the strategy used in Selezyme. Briefly, each molecule in a reaction was represented using a Morgan fingerprint, and pairwise molecular similarity was calculated using the Tanimoto coefficient. For two reactions, Reaction 1 and Reaction 2, we considered both direct matching and cross-matching between reactants and products.

For direct matching, reactants were compared with reactants and products were compared with products:

$$S_{1}=\sqrt{\frac{\mathrm{sim}(R_{1},R_{2})^{2}+\mathrm{sim}(P_{1},P_{2})^{2}}{2}}$$

For cross-matching, reactants were compared with products and products were compared with reactants:

$$S_{2}=\sqrt{\frac{\mathrm{sim}(R_{1},P_{2})^{2}+\mathrm{sim}(P_{1},R_{2})^{2}}{2}}$$

where $R_{1}$and $P_{1}$denote the reactants and products of Reaction 1, respectively, and $R_{2}$and $P_{2}$denote the reactants and products of Reaction 2, respectively. The final reaction similarity score was defined as:

$$S=\max(S_{1},S_{2})$$

This definition accounts for both direct reactant-product correspondence and possible reversed reaction relationships. Reaction similarity was used to quantify the chemical relatedness between reactions and to support benchmark interpretation under different reaction-similarity regimes.

#### **Supplementary Note 8 | RHEA reaction preprocessing and deduplication**

RHEA reactions were preprocessed to generate a standardized reaction library for enzyme–reaction retrieval and large-scale screening. We used the RHEA direction file and reaction SMILES file as input. For each reaction entry, the left-to-right reaction SMILES was used as the representative direction and was also assigned to the corresponding master and bidirectional RHEA entries to maintain consistency with the model training procedure. Polymer reactions without available reaction SMILES were excluded.

Reaction SMILES were then standardized using RDKit. Redundant species, including *, [H+], and [H]*[H], were removed. Molecules that appeared unchanged on both sides of the reaction were also removed based on Morgan fingerprint Tanimoto similarity of 1. Reactions were excluded if they could not be parsed by RDKit, if either side contained more than 150 atoms, if either side lacked carbon-containing molecules, or if either side contained more than three carbon-containing molecules. Remaining molecules were neutralized and converted into canonical SMILES. Reactants and products were sorted to generate a canonical reaction SMILES representation.

To construct a non-redundant reaction library, standardized reactions with identical CANO_RXN_SMILES were subsequently grouped, and all corresponding RHEA identifiers were retained by concatenating them with a pipe separator. This step preserved the mapping between a chemically unique reaction and its original RHEA records while ensuring that each canonical reaction was represented only once in downstream screening. The resulting deduplicated RHEA reaction set was used as the reaction library for EzHit-based retrieval and OMG_Prot50 × RHEA large-scale screening.

### **Supplementary Figures**

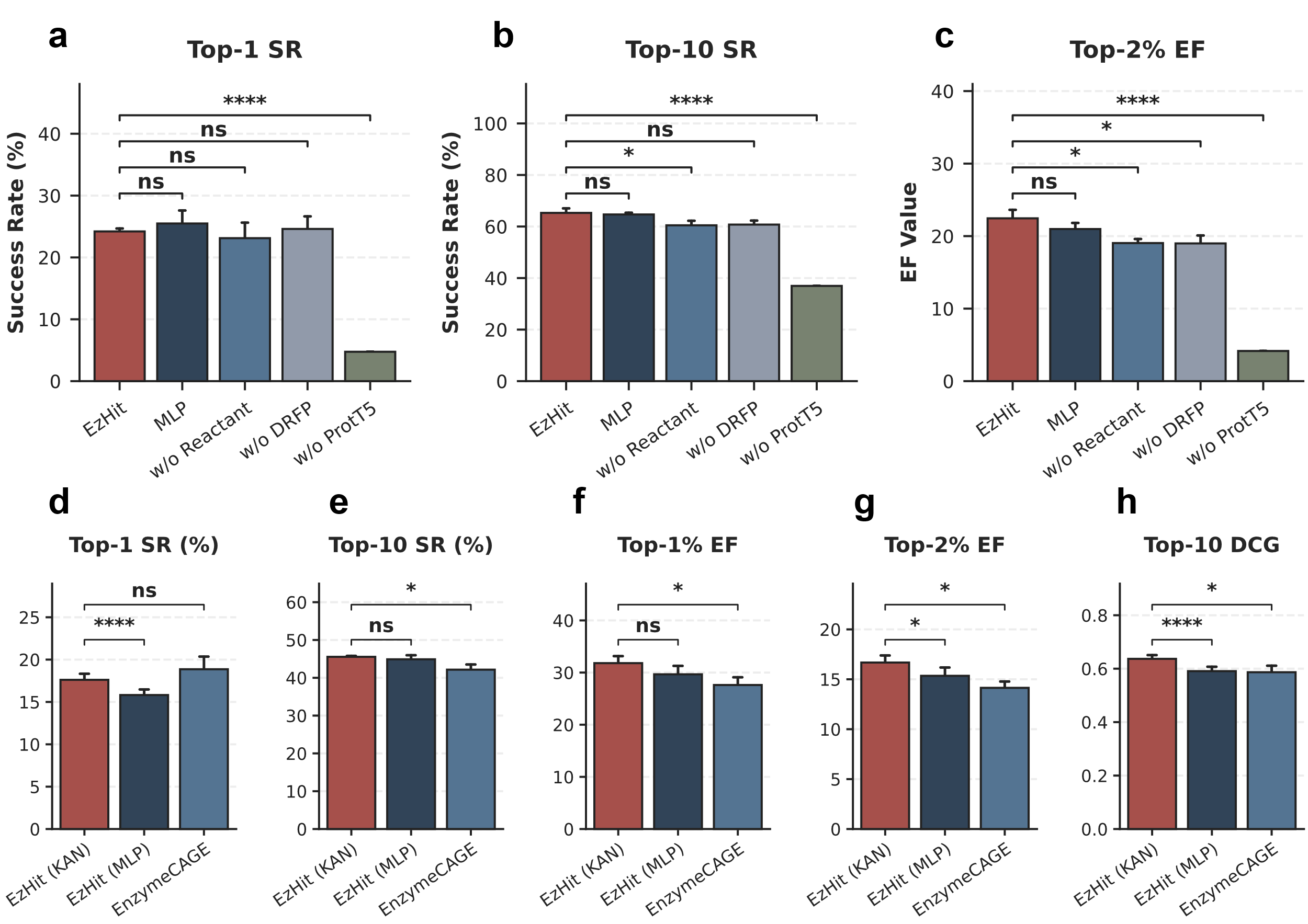

#### **Supplementary Figure 1 | Architectural ablation and comparative evaluation of the EzHit model.**

**a–c**, Ablation study of the EzHit architecture evaluated on the Enzyme-405 dataset. Performance is measured by Top-1 Success Rate (**a**), Top-10 Success Rate (**b**), and Top-2% Enrichment Factor (**c**). The full EzHit model is compared against a variant replacing the KAN module with a standard MLP (MLP), alongside models removing the reactant molecular graph (w/o Reactant), DRFP fingerprint (w/o DRFP), and ProtT5 protein sequence embeddings (w/o ProtT5). **d–h**, Performance comparison on the challenging, highly sparse Orphan-335 dataset to evaluate the specific contribution of the KAN architecture. The KAN-based EzHit model (EzHit (KAN)) is compared with its MLP-based counterpart (EzHit (MLP)) and the state-of-the-art EnzymeCAGE model across Top-1 (**d**) and Top-10 (**e**) Success Rates, Top-1% (**f**) and Top-2% (**g**) Enrichment Factors, and Top-10 DCG (**h**). Across all panels, bar heights represent the mean values from five independent runs with different random seeds, and error bars indicate the standard deviation. Statistical significance was determined using a two-sided paired t-test across five matched random seeds (40-44) comparing the primary EzHit/EzHit (KAN) model against other baselines (ns: not significant, * p < 0.05, ** p < 0.01, *** p < 0.001, **** p < 0.0001).

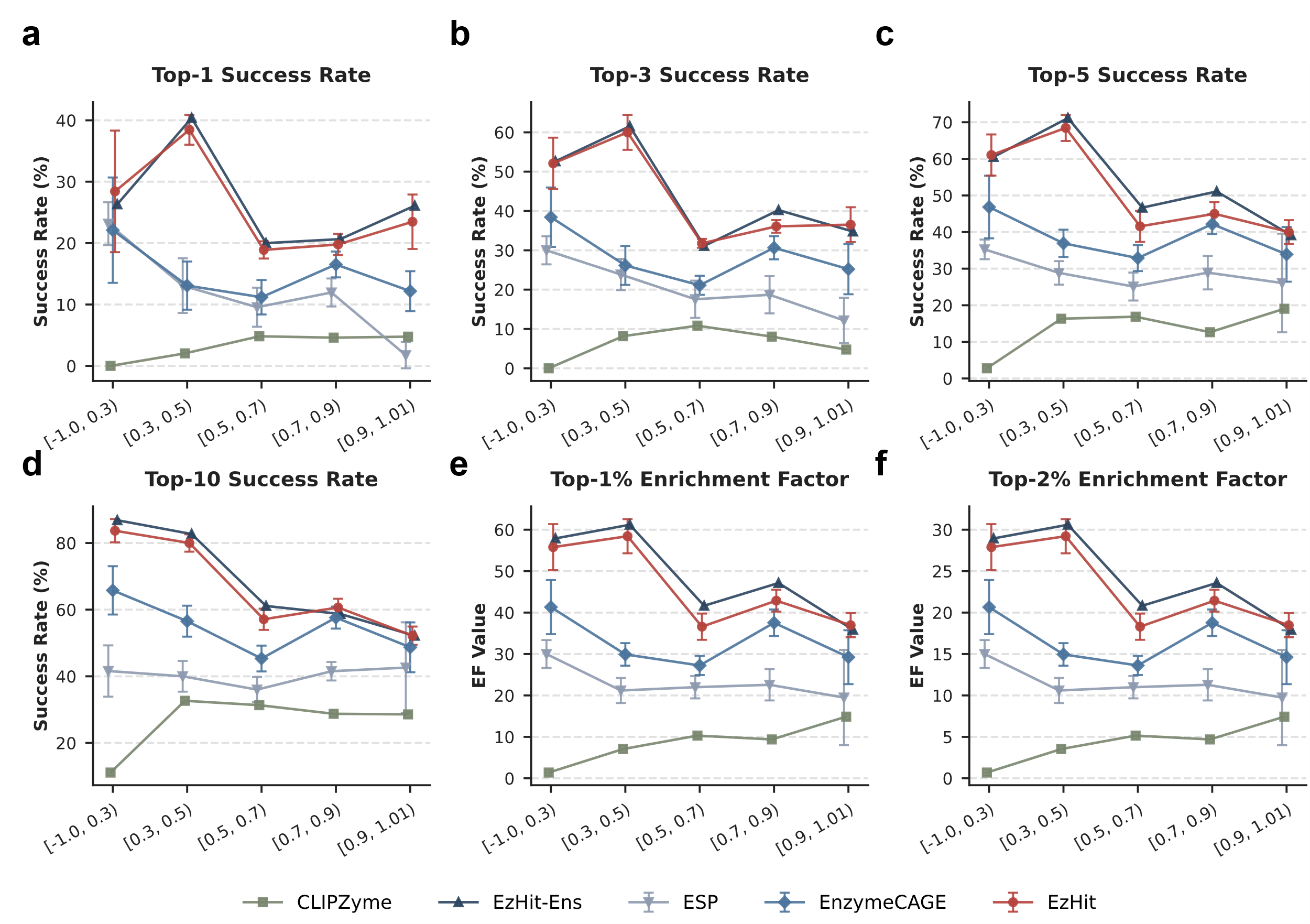

#### **Supplementary Figure 2 | Performance comparison of enzyme-reaction pair retrieval models across different structural similarity cutoffs on the Enzyme-405 dataset.**

Evaluation metrics include Top-1 (**a**), Top-3 (**b**), Top-5 (**c**), and Top-10 (**d**) Success Rates, alongside Top-1% (**e**) and Top-2% (**f**) Enrichment Factors. Test reactions are grouped into five bins based on their maximum structural Tanimoto similarity to the training set. Markers and solid lines denote the mean performance of the evaluated methods (ESP, CLIPZyme, EnzymeCAGE, our proposed single model EzHit, and the ensemble model EzHit-Ens). Error bars represent the standard deviation across five independent runs evaluated under different random seeds (applicable for models with multi-run configurations).

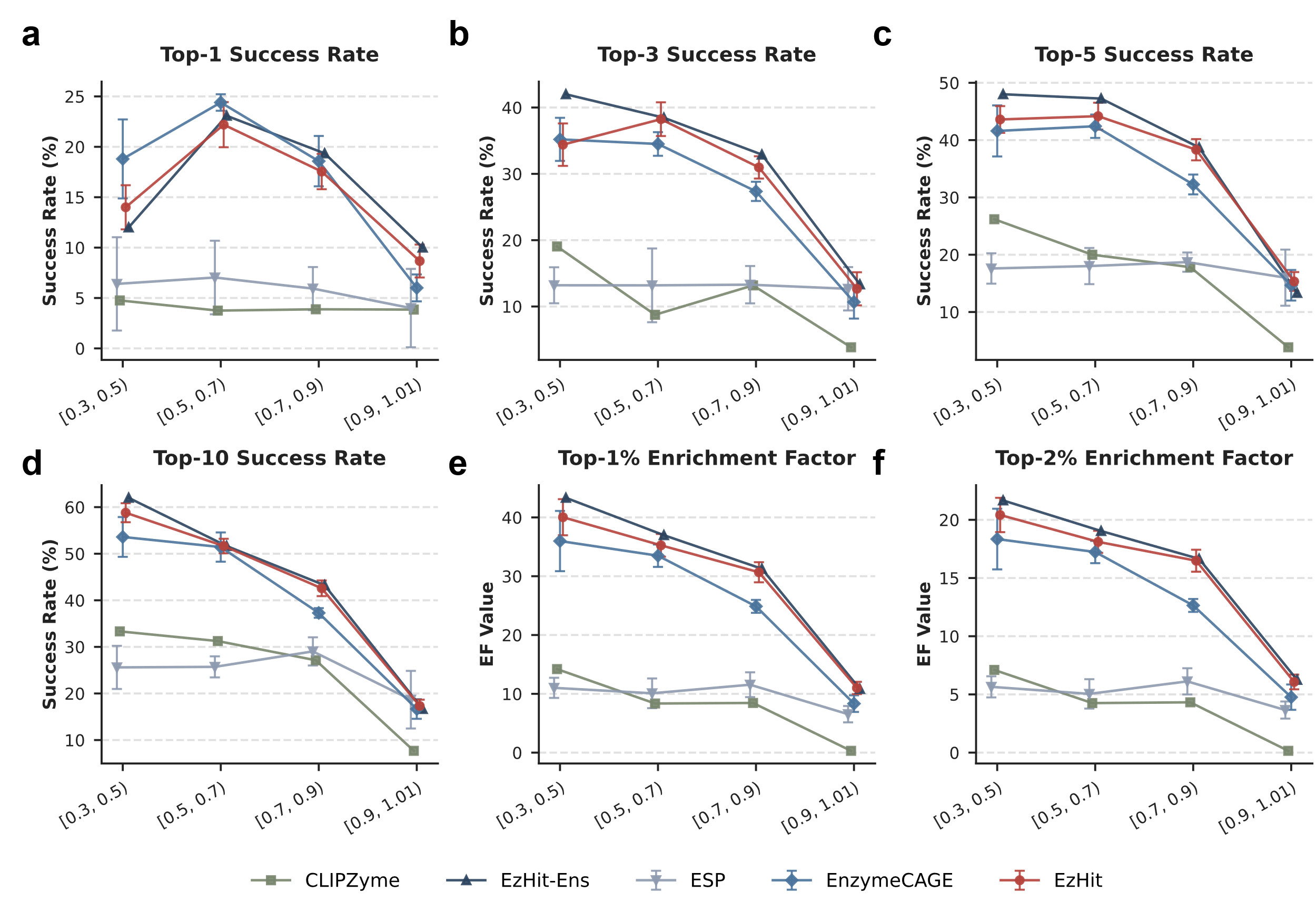

#### **Supplementary Figure 3 | Performance comparison of enzyme-reaction pair retrieval models across different structural similarity cutoffs on the Orphan-335 dataset.**

Evaluation metrics include Top-1 (**a**), Top-3 (**b**), Top-5 (**c**), and Top-10 (**d**) Success Rates, alongside Top-1% (**e**) and Top-2% (**f**) Enrichment Factors. Test reactions are grouped into four bins based on their maximum structural Tanimoto similarity to the training set. Markers and solid lines denote the mean performance of the evaluated methods (ESP, CLIPZyme, EnzymeCAGE, our proposed single model EzHit, and the ensemble model EzHit-Ens). Error bars represent the standard deviation across five independent runs evaluated under different random seeds (applicable for models with multi-run configurations).

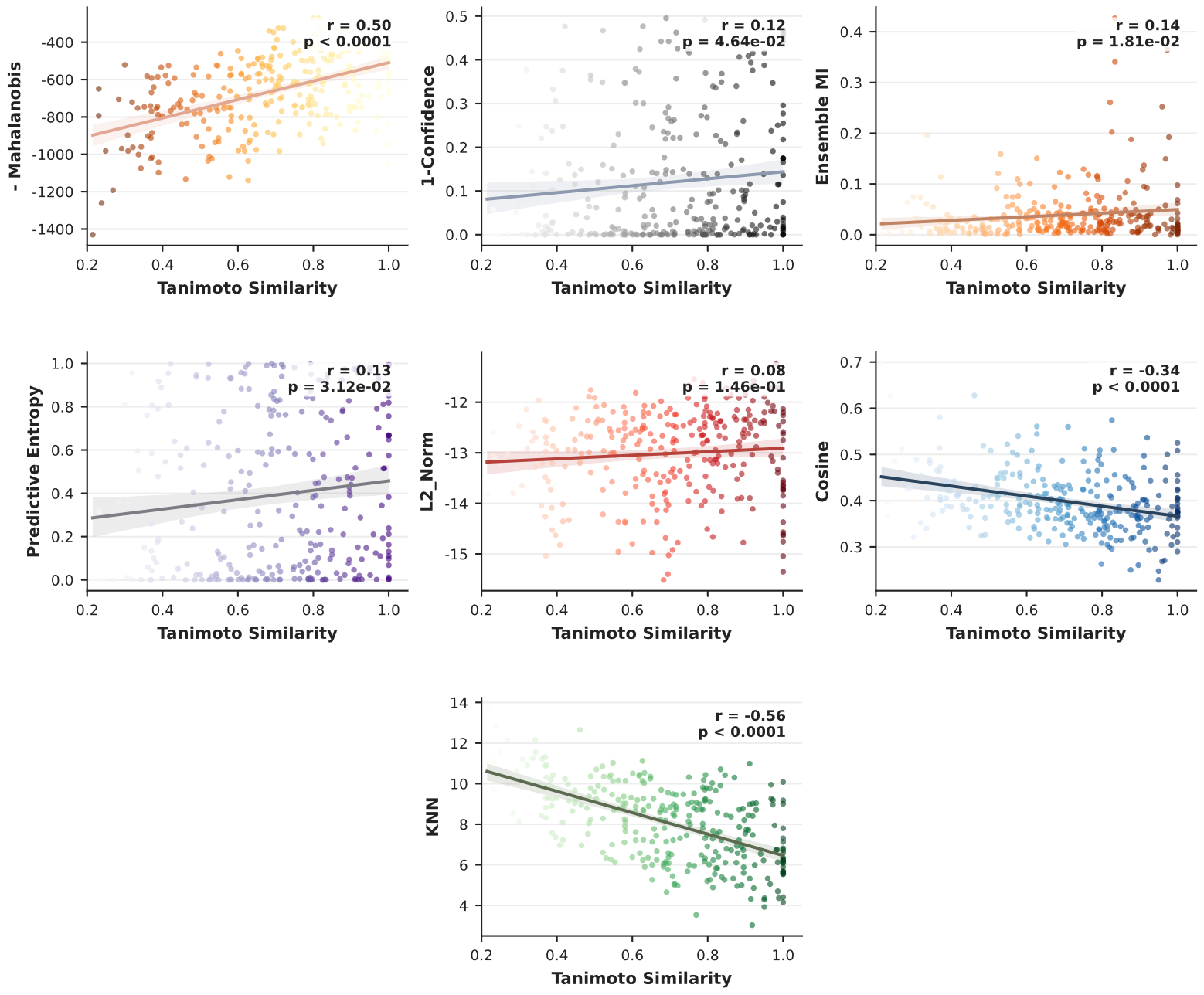

#### **Supplementary Figure 4 | Correlation analysis of all uncertainty quantification (UQ) strategies with structural novelty on the Enzyme-405 dataset.**

Evaluation of seven UQ metrics, including the proposed inverted Mahalanobis distance and six baseline methods, on the in-distribution Enzyme-405 dataset. The scatter plots illustrate the relationship between each UQ score and the maximum structural Tanimoto Similarity to the training set. Solid lines represent linear regression fits, with Pearson’s correlation coefficients (*r*) and associated *p*-values annotated for each UQ strategy.

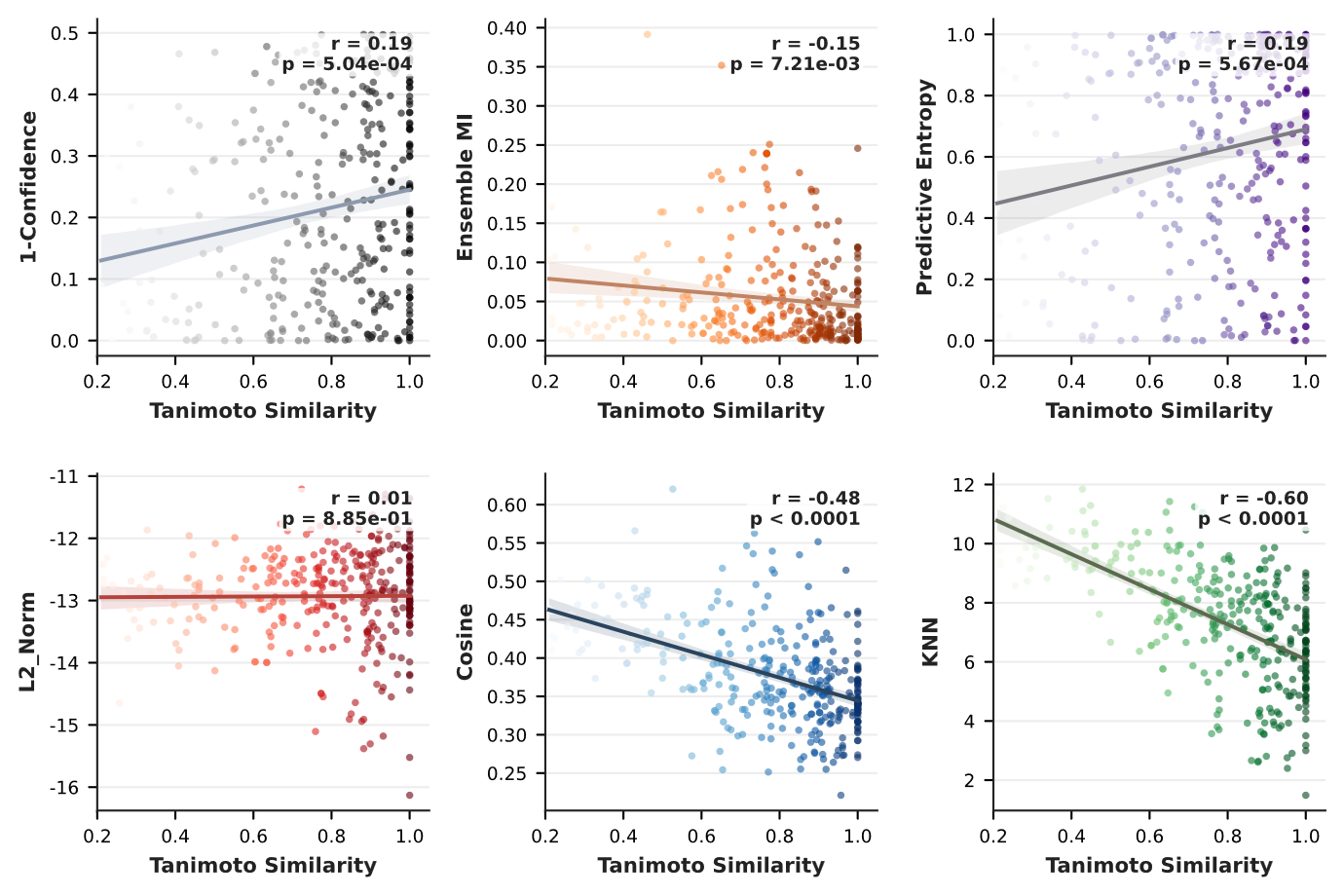

#### **Supplementary Figure 5 | Correlation analysis of alternative uncertainty quantification (UQ) strategies with structural novelty on the Orphan-335 dataset.**

Evaluation of six baseline and geometric UQ metrics on the out-of-distribution (OOD) Orphan-335 dataset. The scatter plots illustrate the relationship between each UQ score and the maximum structural Tanimoto Similarity to the training set. Solid lines represent linear regression fits, with Pearson’s correlation coefficients (*r*) and associated *p*-values annotated for each UQ strategy.

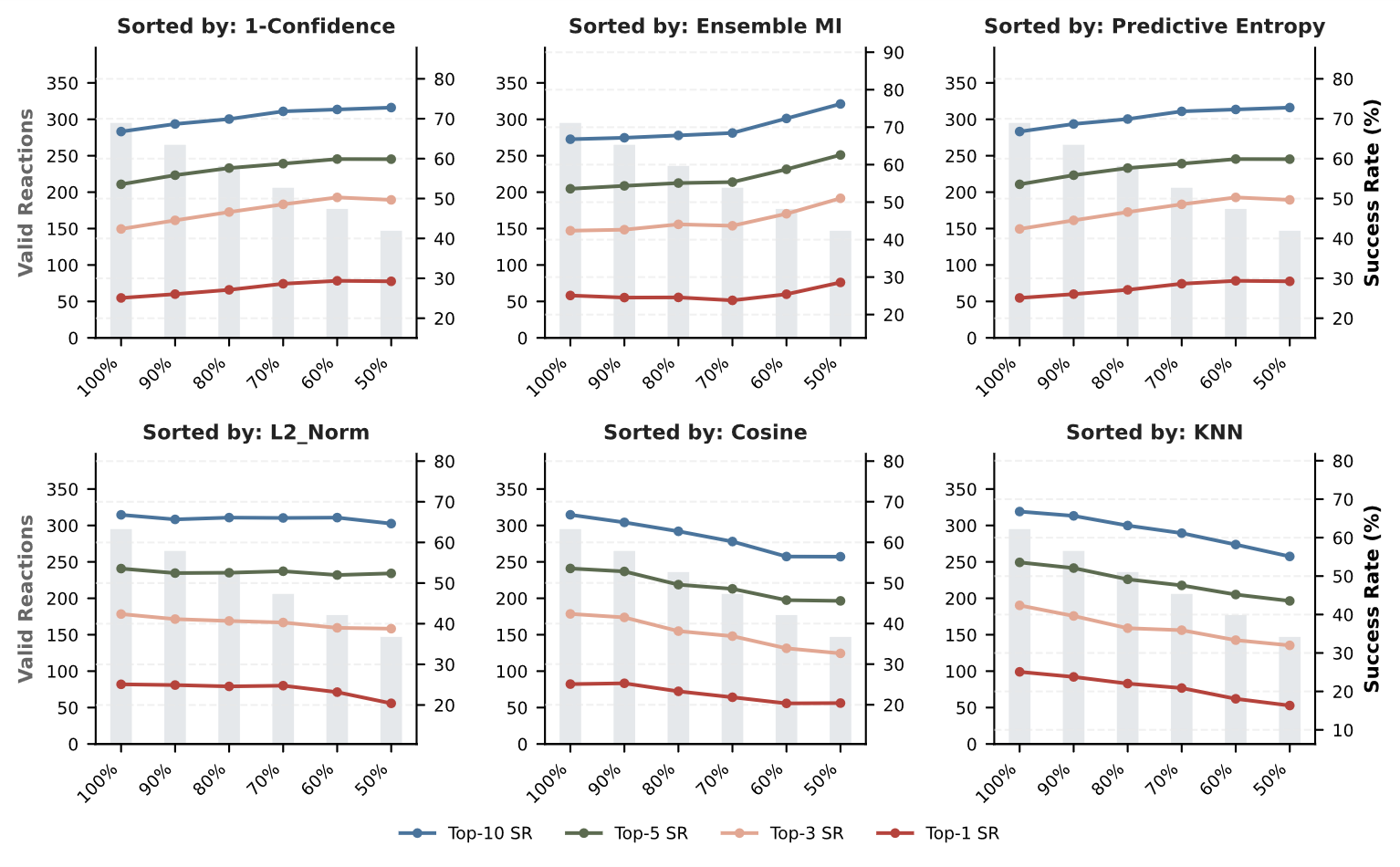

#### **Supplementary Figure 6 | Confidence-based retention curves using alternative UQ strategies for the Enzyme-405 dataset.**

Confidence-based rejection curves for the Enzyme-405 dataset utilizing six alternative UQ metrics as filtering criteria. The line plots (right y-axis) track the Top-1 to Top-10 Success Rates as reactions are progressively filtered from 100% down to 50% retention based on the respective UQ scores. The bar plots (left y-axis) indicate theabsolute number of retained valid reactions at each retention threshold.

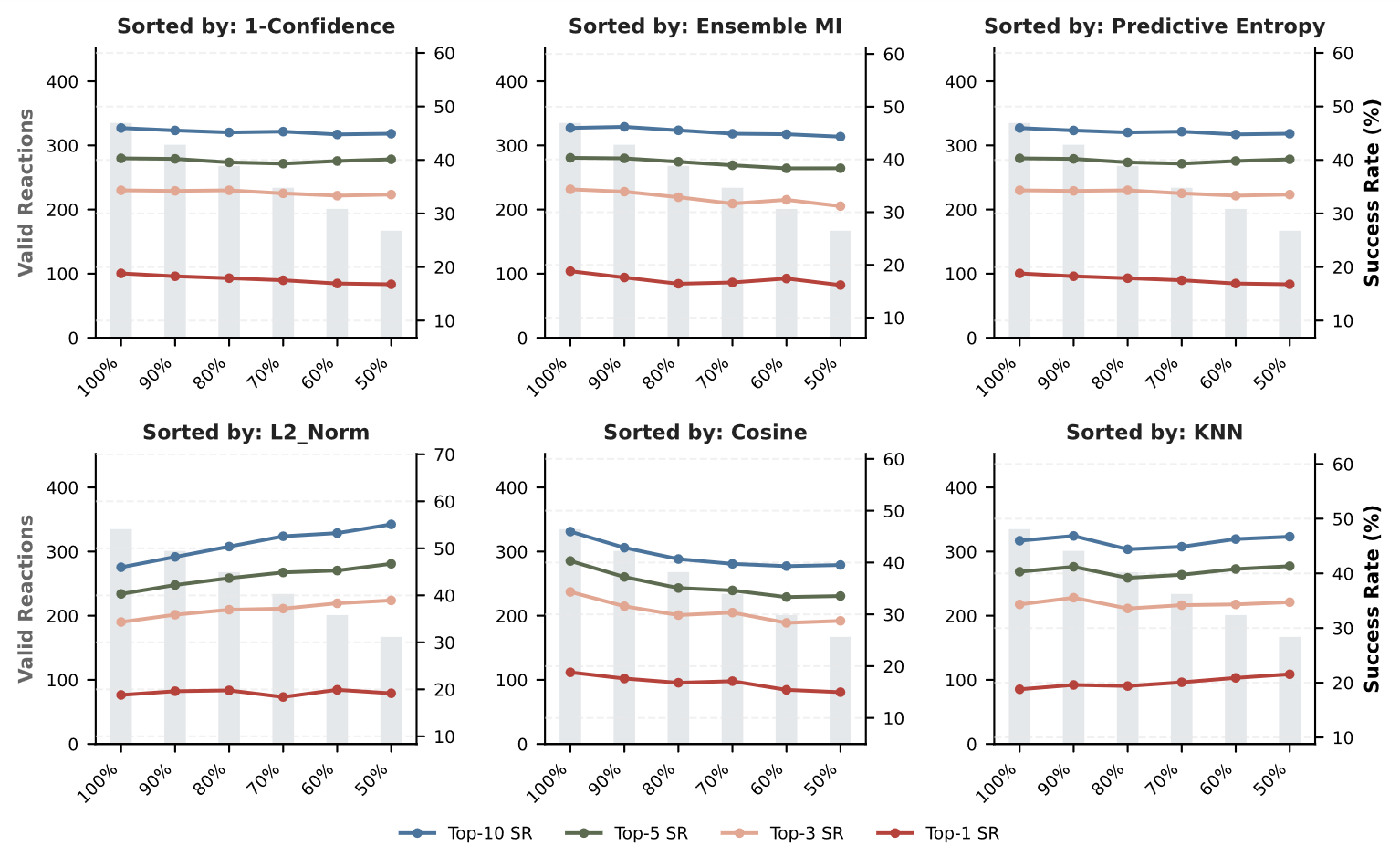

#### **Supplementary Figure 7 | Confidence-based retention curves using alternative UQ strategies for the Orphan-335 dataset.**

Confidence-based rejection curves for the Orphan-335 dataset utilizing six alternative UQ metrics as filtering criteria. The line plots (right y-axis) track the Top-1 to Top-10 Success Rates as reactions are progressively filtered from 100% down to 50% retention based on the respective UQ scores. The bar plots (left y-axis) indicate the absolute number of retained valid reactions at each retention threshold.

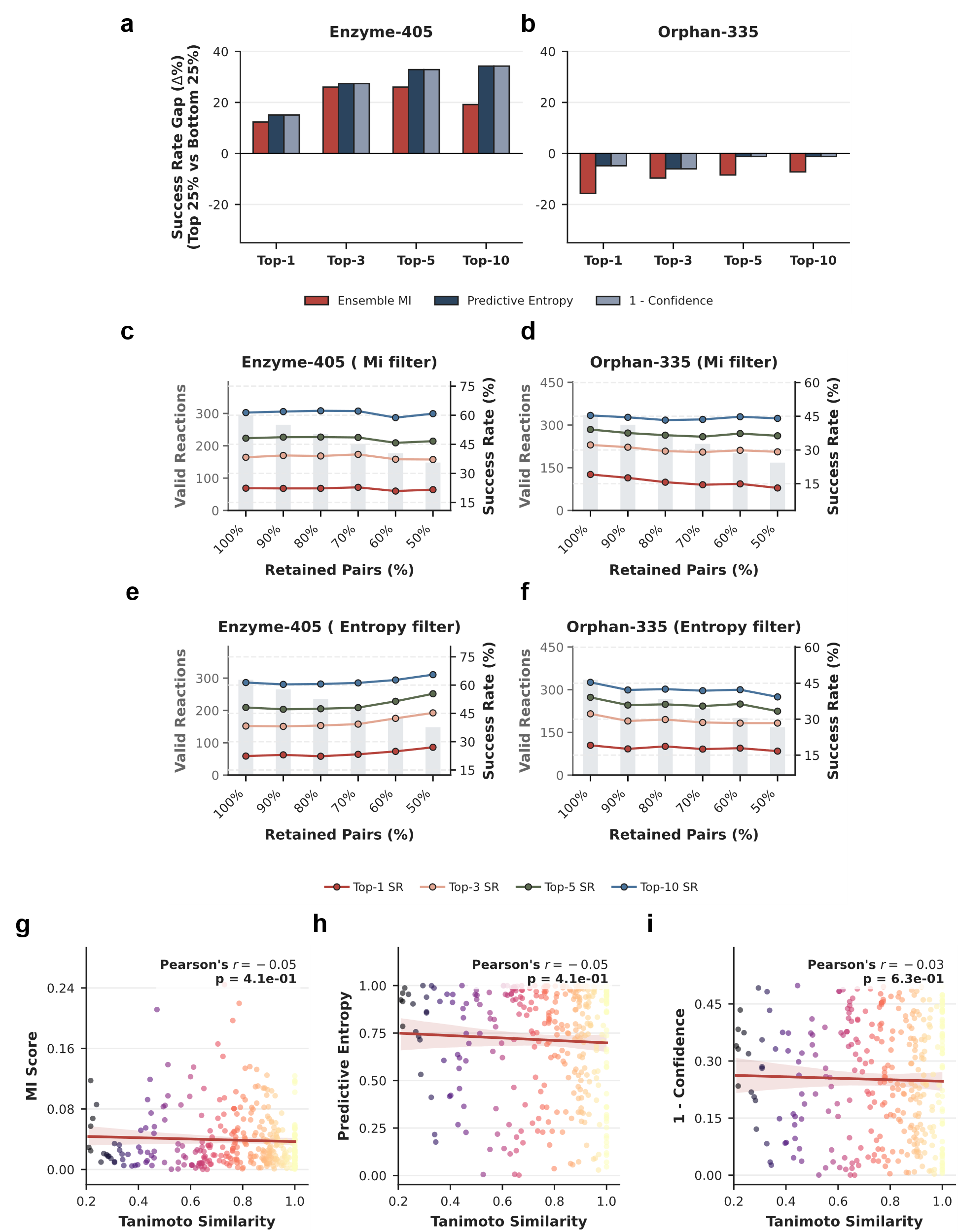

#### **Supplementary Figure 8 | Evaluation of uncertainty quantification metrics using a 5-fold cross-validation ensemble.**

**a, b**, Evaluation of uncertainty metric calibration, measured by the Success Rate Gap (Δ%) between the top 25% most confident and bottom 25% least confident predictions. Performance comparisons among Ensemble MI, Predictive Entropy, and 1 - Confidence are shown across Top-1, Top-3, Top-5, and Top-10 rankings on the Enzyme-405 (**a**) and Orphan-335 (**b**) test sets. **c–f**, Retention curves illustrating the model Success Rate (line plots, right y-axis) and the number of retained valid reactions (bar plots, left y-axis) when progressively filtered from 100% down to 50% based on prediction confidence. Filtering is executed using the Ensemble MI score for the Enzyme-405 (**c**) and Orphan-335 (**d**) datasets, and using the Predictive Entropy score for the Enzyme-405 (**e**) and Orphan-335 (**f**) datasets. **g–i**, Correlation between individual uncertainty metrics and the maximum structural Tanimoto Similarity to the training set for reactions in the Orphan-335 dataset. The metrics evaluated include Ensemble MI Score (**g**), Predictive Entropy (**h**), and 1 - Confidence (**i**). Solid lines represent linear regression fits, with Pearson’s correlation coefficients (*r*) and the associated *p*-values annotated.

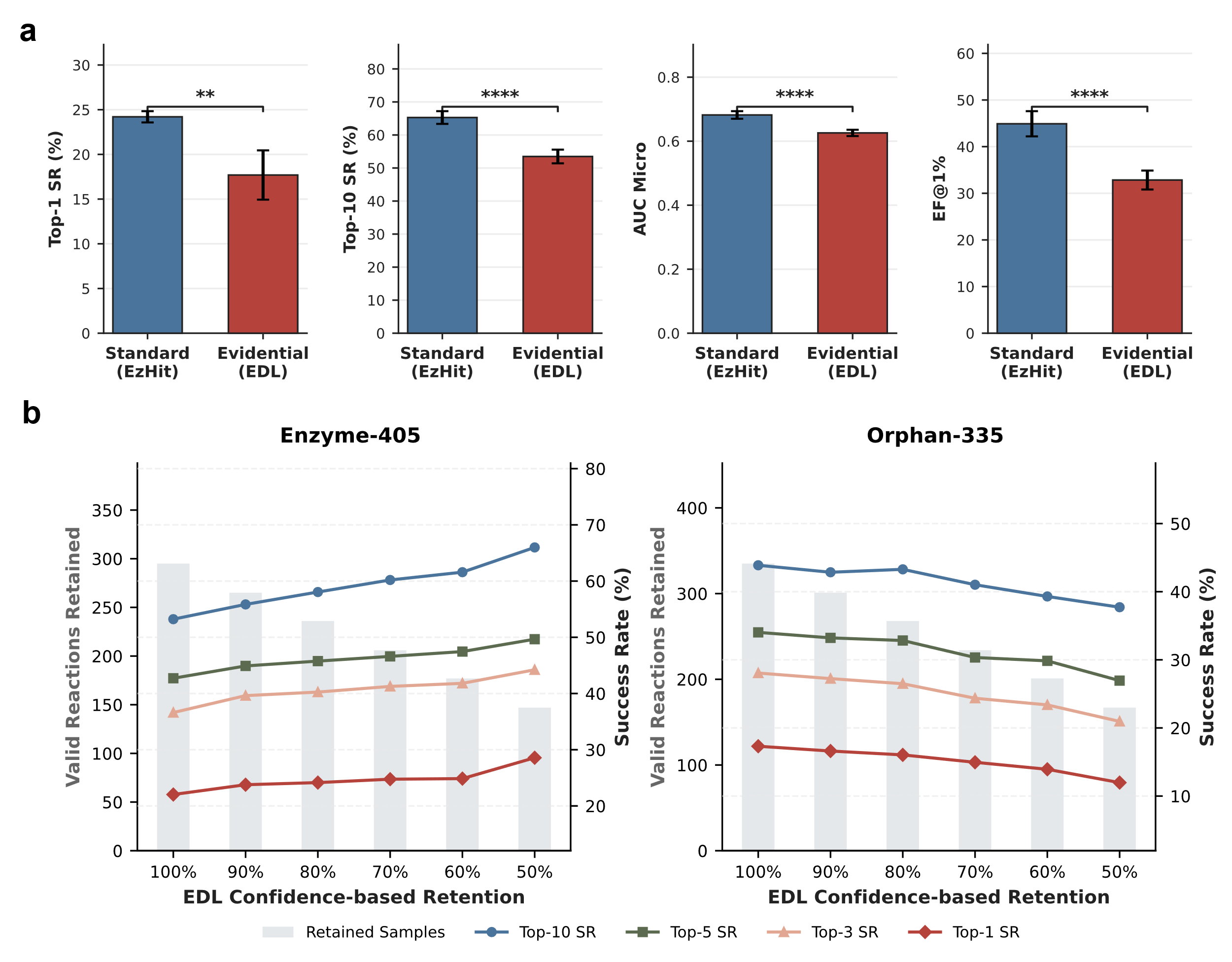

#### **Supplementary Figure 9 | Predictive performance degradation and out-of-distribution calibration failure induced by Evidential Deep Learning (EDL) regularization.**

**a,** Comparison of the standard KAN model (EzHit) and the evidential KAN (trained with EDL loss) on the Enzyme-405 dataset. The bar plots display the mean performance across four evaluation metrics, namely Top-1 Success Rate, Top-10 Success Rate, AUC Micro, and Enrichment Factor at 1% (EF@1%). Data are presented as mean ± s.d. derived from independent training runs with five different random initializations (*n* = 5). Statistical significance was determined using an independent two-sided Student’s *t*-test ( *P* < 0.01, **** *P* < 0.0001). **b,** Confidence-based retention curves utilizing the epistemic uncertainty derived from the EDL framework. Results are shown for the in-distribution Enzyme-405 dataset (left) and the out-of-distribution Orphan-335 dataset (right). The left y-axis (gray bars) represents the absolute number of valid reactions retained at varying confidence thresholds. The right y-axis (colored lines) traces the corresponding top-*k* success rates.

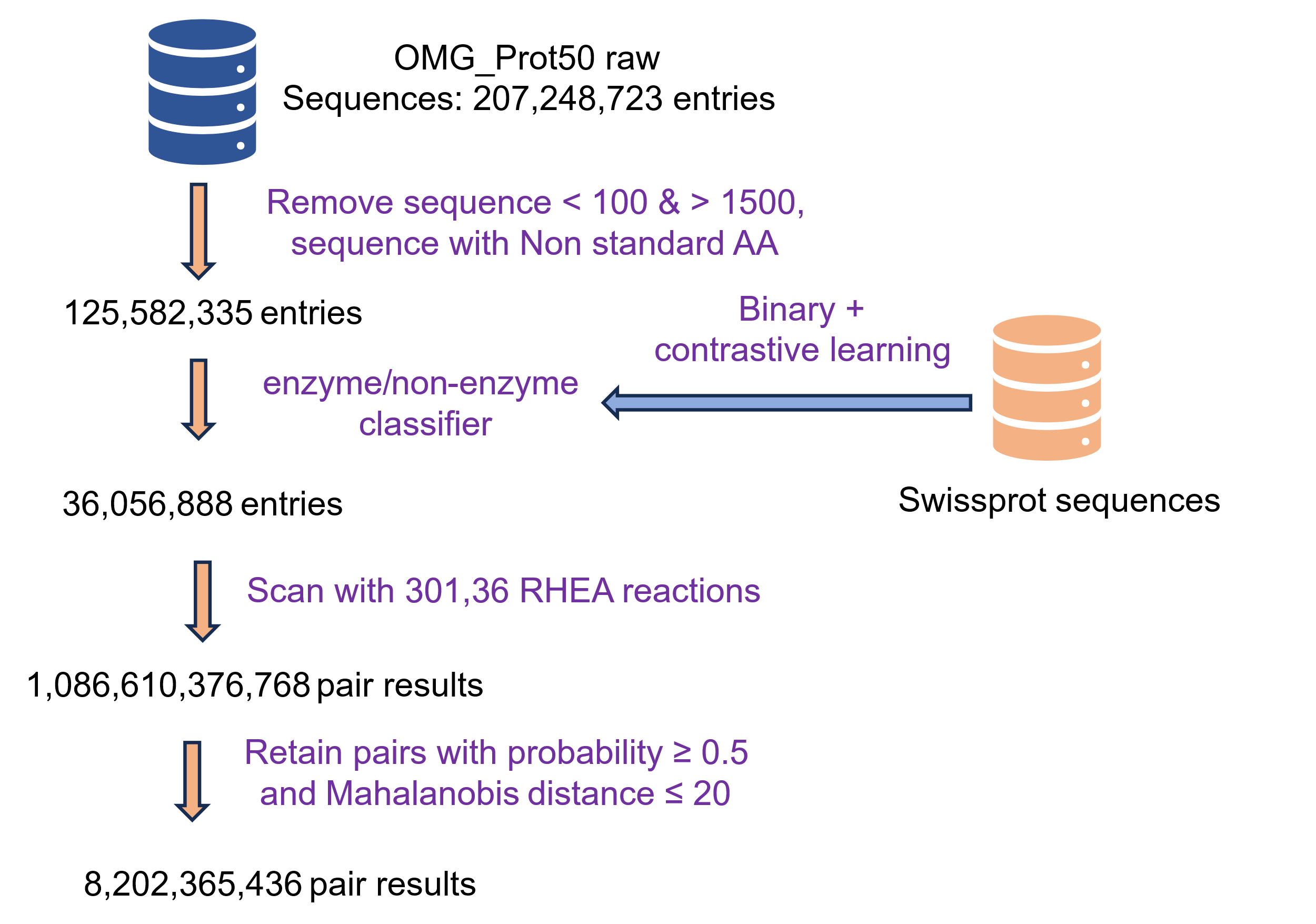

#### **Supplementary Figure 10. Overview of the OMG_Prot50-scale enzyme–reaction screening workflow.**

Raw OMG_Prot50 sequences were cleaned by length and amino-acid-composition filters, followed by enzyme/non-enzyme pre-filtering using a Swiss-Prot-trained classifier. The retained putative enzymes were screened against 30,136 RHEA reactions using EzHit. Prediction pairs satisfying the probability and Mahalanobis-distance thresholds were retained for downstream large-scale analysis.

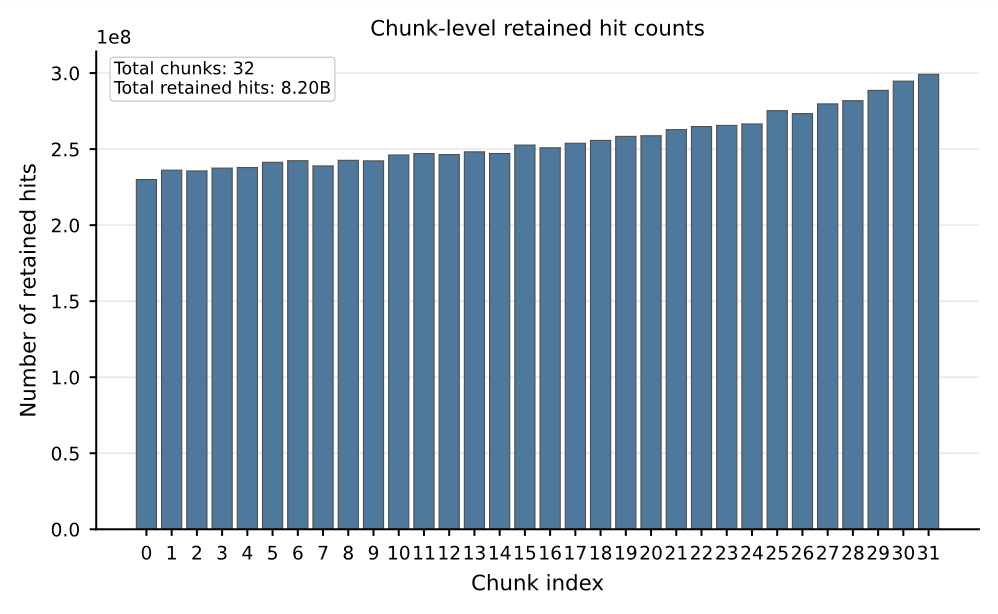

#### **Supplementary Figure 11 |. Chunk-level retained hit counts across the OMG_Prot50 screening run.**

Each bar represents one of the 32 processed OMG_Prot50 chunks, and the bar height indicates the number of retained EzHit enzyme–reaction hits in that chunk. Retained hits were defined as enzyme–reaction pairs with predicted probability > 0.5 and Mahalanobis distance < 20. The relatively consistent chunk-level hit counts indicate that the large-scale screening procedure was completed across all protein chunks without obvious chunk-specific failures.

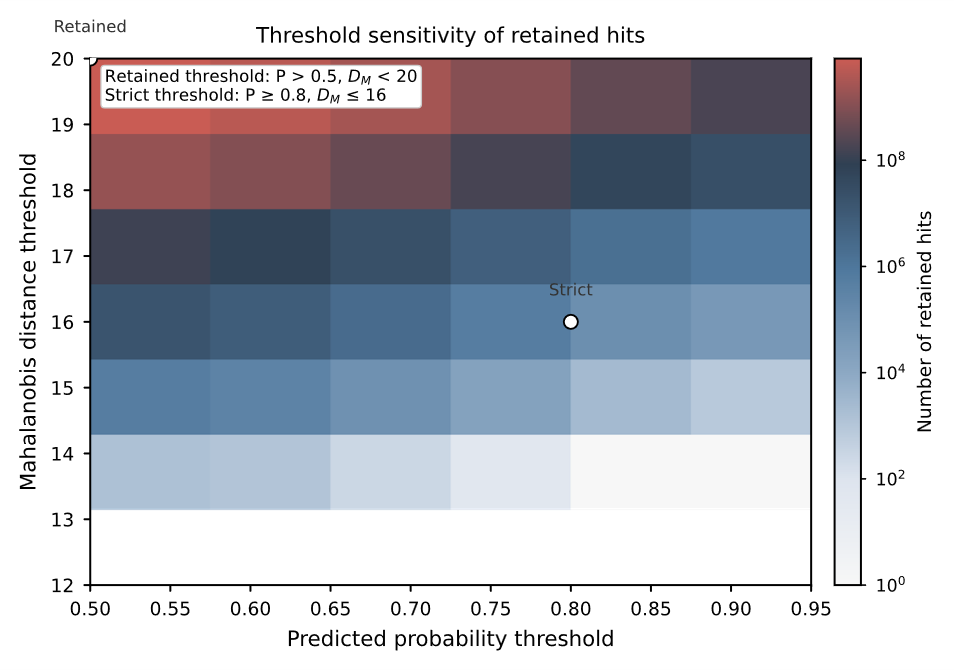

#### **Supplementary Figure 12 | Sensitivity of retained hit counts to probability and Mahalanobis-distance thresholds.**

The heatmap shows the number of retained enzyme–reaction hits obtained under different combinations of predicted probability thresholds and Mahalanobis-distance thresholds. The retained-hit cutoff used for the global association map was defined as predicted probability > 0.5 and Mahalanobis distance < 20, whereas the stricter cutoff used for high-confidence downstream analyses was defined as predicted probability ≥ 0.8 and Mahalanobis distance ≤ 16. The gradual change in retained hit counts across threshold combinations supports the use of probability and Mahalanobis distance as complementary filters for confidence-controlled large-scale enzyme–reaction matching.

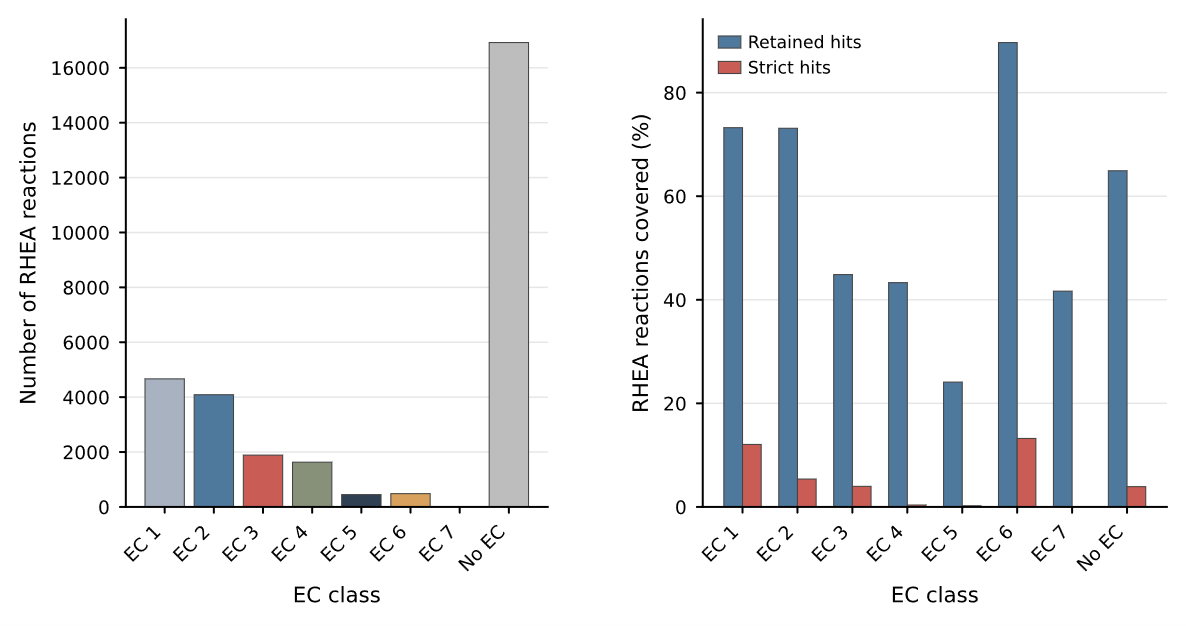

#### **Supplementary Figure 13 |. EC-class and No-EC coverage of RHEA reactions by EzHit.**

The left panel shows the number of RHEA reactions in each EC class and in the No-EC group. The right panel shows the percentage of reactions in each group covered by retained hits and strict hits. Retained hits were defined as predicted probability > 0.5 and Mahalanobis distance < 20, and strict hits were defined as predicted probability ≥ 0.8 and Mahalanobis distance ≤ 16.

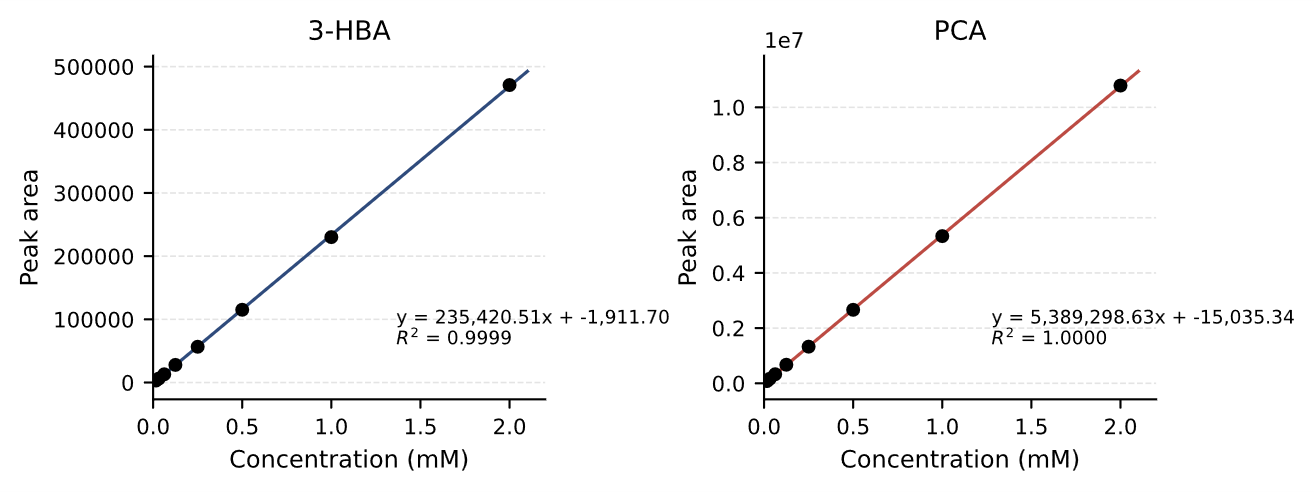

#### **Supplementary Figure 14 | Standard curves of 3-hydroxybenzoate and protocatechuate used for HPLC-based quantification.**

Calibration curves were generated from authentic 3-hydroxybenzoate and protocatechuate standards and used for quantification of substrate conversion and product formation, respectively.

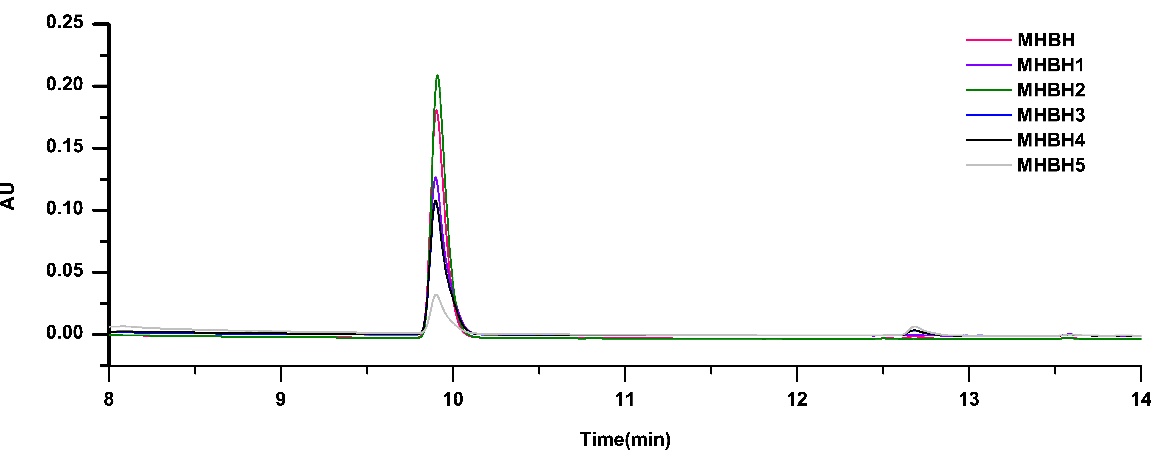

#### **Supplementary Figure 15 | HPLC analysis of MHBH and EzHit-prioritized metagenomic candidates for RHEA:11480.**

Overlaid chromatograms are shown for the characterized MHBH reference and five EzHit-prioritized candidates, MHBH1–MHBH5, following incubation with 3-hydroxybenzoate and NADPH. Formation of protocatechuate was identified by comparison with an authentic standard, and all samples were analyzed under the same chromatographic conditions

### **Supplementary Tables**

#### **Supplementary Table 1. Performance Metrics on Enzyme405 test set**

| **Model** | **Top10DCG** | **SR1(%)** | **SR10(%)** | **EF0.01** | **AUC** |
| --- | --- | --- | --- | --- | --- |
| EzHit-Ens | 0.5875 | 25.08 | 66.78 | 48.45 | 0.6932 |
| EzHit | 0.5708 | 24.41 | 65.42 | 46.53 | 0.6988 |
| EnzymeCAGE | 0.4307 | 17.29 | 54.24 | 34.39 | 0.6338 |
| ESP | 0.3464 | 14.24 | 40.34 | 21.02 | 0.5718 |
| CLIPzyme | 0.1545 | 3.62 | 30.43 | 10.04 | 0.4904 |

#### **Supplementary Table 2. Performance Metrics on Orphan335 test set**

| **Model** | **Top10DCG** | **SR1(%)** | **SR10(%)** | **EF0.01** | **AUC** |
| --- | --- | --- | --- | --- | --- |
| EzHit-Ens | 0.6637 | 18.81 | 45.97 | 33.04 | 0.8055 |
| EzHit | 0.6503 | 18.81 | 45.97 | 33.96 | 0.8041 |
| EnzymeCAGE | 0.5781 | 18.21 | 42.39 | 26.86 | 0.6090 |
| ESP | 0.4225 | 9.01 | 42.49 | 16.59 | 0.6739 |
| CLIPzyme | 0.2936 | 3.87 | 28.87 | 8.86 | 0.5368 |

#### **Supplementary Table 3. Performance Metrics on External test set**

| **Test set** | **Model** | **SR1(%)** | **SR3(%)** | **SR5(%)** |
| --- | --- | --- | --- | --- |
| P450 | EzHit-ft-Ens | 13.6 | 26.7 | 33.5 |
|  | EzHit-ft | 12.6 | 24.6 | 31.4 |
|  | EzHit-Ens | 2.6 | 13.1 | 19.9 |
|  | EzHit | 2.6 | 10.5 | 19.9 |
|  | EnzymeCAGE-ft | 10.5 | 16.2 | 22.0 |
|  | EnzymeCAGE | 7.3 | 12.0 | 15.2 |
| Phosphatase | EzHit-ft-Ens | 18.5 | 50.3 | 63.6 |
|  | EzHit-ft | 19.9 | 51.0 | 60.9 |
|  | EzHit-Ens | 26.5 | 57.0 | 66.9 |
|  | EzHit | 22.5 | 53.0 | 68.2 |
|  | EnzymeCAGE-ft | 21.2 | 47.7 | 56.3 |
|  | EnzymeCAGE | 9.3 | 39.1 | 56.3 |
| Terpene | EzHit-ft-Ens | 3.3 | 9.8 | 17.4 |
|  | EzHit-ft | 6.5 | 15.2 | 22.8 |
|  | EzHit-Ens | 4.3 | 10.9 | 18.5 |
|  | EzHit | 5.4 | 14.1 | 18.5 |
|  | EnzymeCAGE-ft | 10.9 | 17.4 | 30.4 |
|  | EnzymeCAGE | 6.5 | 10.9 | 22.8 |

#### **Supplementary Table 4. Hyperparameters used for training the enzyme/non-enzyme pre-filtering classifier**

| **Hyperparameter** | **Value** | **Description** |
| --- | --- | --- |
| input_dim | 1024 | Dimension of ProtT5 sequence embedding |
| encoder_dim | 512 | Dimension of shared hidden representation |
| projection_dim | 128 | Dimension of contrastive projection head |
| classifier_hidden_dim | 128 | Hidden dimension of binary classification head |
| activation | GELU | Activation function used in hidden layers |
| temperature | 0.1 | Temperature parameter for supervised contrastive loss |
| α | 0.3 | Weight of supervised contrastive loss |
| 1-α | 0.7 | Weight of weighted binary cross-entropy loss |
| train/validation split | 8:2 | Ratio of training and validation sets |
| split strategy | stratified | Preserves enzyme/non-enzyme label distribution |
| batch_size | 4096 | Number of samples per training batch |
| n_epochs | 50 | Maximum number of training epochs |
| learning_rate | 0.001 | Initial learning rate for AdamW |
| weight_decay | 0.0001 | Weight decay in AdamW |
| early_stopping_patience | 5 | Early stopping patience |
| min_delta | 0.0001 | Minimum improvement required for checkpoint update |
| probability_threshold | 0.5 | Threshold for retaining putative enzyme sequences |

#### **Supplementary Table 5. Hyperparameters used for training EDL-KAN uncertainty**

| **Hyperparameter** | **Value** | **Description** |
| --- | --- | --- |
| fp_dim | 2048 | Dimension of DRFP reaction fingerprint |
| reactant_fp_dim | 2048 | Dimension of reactant Morgan fingerprint |
| hidden | 512 | Hidden dimension of the compressor and KAN module |
| output_dim | 2 | Evidence values for binary classification |
| evidence_activation | Softplus | Ensures non-negative evidence output |
| λ_EDL_ | 0.1 | Weight of KL divergence in EDL loss |
| λ_KAN_ | 0.0001 | Weight of KAN regularization loss |
| batch_size | 2048 | Number of samples per training batch |
| n_epochs | 30 | Maximum number of training epochs |
| learning_rate | 0.001 | Initial learning rate for AdamW |
| weight_decay | 0.0001 | Weight decay in AdamW |
| gradient_clip_norm | 1.0 | Maximum gradient norm |
| select_metric | Top-10 Success Rate | Validation metric used for checkpoint selection |
| patience | 5 | Early stopping patience |
| min_delta | 0.003 | Minimum improvement required for checkpoint update |
| warmup_epochs | 2 | Minimum epochs before early stopping |
| uq_threshold | 0.4 | Threshold for EDL uncertainty-based filtering |
| random_seed | 42 | Default random seed |

#### **Supplementary Table 6. Hyperparameters of the standard EzHit KAN model**

| **Hyperparameter** | **Value** | **Description** |
| --- | --- | --- |
| fp_dim | 2048 | Dimension of DRFP reaction fingerprint |
| reactant_fp_dim | 2048 | Dimension of reactant Morgan fingerprint |
| output_dim | 1 | Single logit for binary classification |
| hidden | 512 | Hidden dimension of the compressor and KAN module |
| dropout | 0.4 | Dropout rate in the compressor module |
| loss_function | weighted BCEWithLogitsLoss | Binary classification loss with class weighting |
| pos_weight | auto | Computed as *N*_negative_ / *N*_positive_ |
| optimizer | AdamW | Optimizer used for model training |
| learning_rate | 1e-4 | Initial learning rate |
| weight_decay | 0.0001 | Weight decay used in AdamW |
| batch_size | 2048 | Number of samples per training batch |
| n_epochs | 30 | Maximum number of training epochs |
| kan_reg_weight | 0.0001 | Weight of KAN regularization loss |
| gradient_clip_norm | 1.0 | Maximum gradient norm |
| select_metric | Top 10 Success Rate | Validation metric for checkpoint selection |
| patience | 5 | Early stopping patience |
| min_delta | 0.003 | Minimum improvement required for checkpoint update |
| warmup_epochs | 2 | Minimum epochs before early stopping |
| random_seed | 42 | Default random seed |

#### **Supplementary Table 7. Hyperparameters of the standard EzHit KAN model**

| **Model** | **Runtime definition** | **Checkpoint** | **Batch size** | **Input pairs** | **Time (s)** | **Throughput (pairs/s)** | **Seconds per 1M pairs** |
| --- | --- | --- | --- | --- | --- | --- | --- |
| EzHit | Cached-feature inference | 42 | 256 | 15,921 | 0.101 ±0.003 | 157,696 | 6.34 |
| EzHit | Cached-feature inference | 42 | 8192 | 15,921 | 0.0749 ±0.0003 | 212,631 | 4.70 |
| EnzymeCAGE | DataLoader + official model.evaluate inference | 42 | 256 | 15,611 | 39.36 ± 0.97 | 397 | 2521.00 |

#### **Supplementary Table 8. Protein sequences used for experimental validation of RHEA:11480.**

| Enzyme | Sequence ID | Sequence identity to MHBH (%) | Protein sequences |
| --- | --- | --- | --- |
| MHBH | - | 100 | MQFHLNGFRPGNPLIAPASPLAPAHTEAVPSQVDVLIVGCGPAGLTLAAQLAAFPDIRTCIVEQKEGPMELGQADGIACRTMEMFEAFEFADSILKEACWINDVTFWKPDPAQPGRIARHGRVQDTEDGLSEFPHVILNQARVHDHYLERMRNSPSRLEPHYARRVLDVKIDHGAADYPVTVTLERCDAAHAGQIETVQARYVVGCDGARSNVRRAIGRQLVGDSANQAWGVMDVLAVTDFPDVRYKVAIQSEQGNVLIIPREGGHLVRFYVEMDKLDADERVASRNITVEQLIATAQRVLHPYKLDVKNVPWWSVYEIGQRICAKYDDVADAVATPDSPLPRVFIAGDACHTHSPKAGQGMNFSMQDSFNLGWKLAAVLRKQCAPELLHTYSSERQVVAQQLIDFDREWAKMFSDPAKEGGQGGVDPKEFQKYFEQHGRFTAGVGTHYAPSLLTGQASHQALASGFTVGMRFHSAPVVRVSDAKPLQLGHCGKADGRWRLYAFAGQNDLAQPESGLLALCRFLESDAASPLRRFTPSGQDIDSIFDLRAIFPQAYTEVALETLPALLLPPKGQLGMIDYEKVFSPDLKNAGQDIFELRGIDRQQGALVVVRPDQYVAQVLPLGDHAALSAYFESFMRA |
| MHBH1 | 3300046814\|Ga0495514_0000026\|CDS\|Ga0495514_0000026_4317_6188\|+\|4317:6188 | 46.3 | MQDAAGTGIDRPGELPDEMDVLVVGSGPAGMLLAAQLSQFPDVTTRIIERRDGRLVLGQADGIQPRSVETFQAFGFAERIVAEAYNIAYMNFWGPDPENPGSIRRTARTEDYAFKISEFPHLIVNQARVLDYFAEAAAHGPGRIVPDYGIEFLGLTVRDSADPHSGRHPVEVRVRYTAGERAGEERTVRATYVVGCDGARSGVREAIGRKHLGGISAHAWGVMDVLVSTDFPDWRTKCAINAEAGNILHIPREGGYLSRMYIDLGAVSDDDGHLVRQTPIDEIIRKANEILHPYTLDVKQVAWHSVYEVGHRVTDRFDDGGEGRIPHVFLTGDACHTHSAKAGQGMNVSMQDGFNLGWKLGHVLTGLAPAELLETYSAERQPVAQQLIDFDREWSSLMARKPEEITDPQELATFYLGTAEFPSGFMTRYGPSQIVTDATHQPLAGGFPIGKRFTSVEVTRVCDGNVIHLGHHAKADGRWRIYAFADAAAGEGSALASWAEWLGSSSSPIVRHTPAGADVDAVFDVKAIYQQRFEDVDISRVPKAFLPTSGPLGLTDWEKVYAAGPDGWRTVDIFDERELSRDGVVVVVRPDQYVAAVLPLTATDELSAFFESALLPQHVVATT |
| MHBH2 | 3300009784\|Ga0123357_10000120\|CDS\|Ga0123357_1000012042\|-\|46403:48292 | 45.3 | MQFHDQGYVSADPRVRPAAGVGLRCSDAFPDNVDVLIIGTGPAGIITAAQLSQFPSIDTRIIERRSGRLELGHADGVNARSVESFAAFGFADEIIAEAYTNNQTSFWRPDPKDPSRIFRASRTADDPTGISEFPHILVNQARVADYFIDYMAKSPARLKPDYGWEFVDLTIDDAQEYPVTVTIRRSAYPQEGVEKIVKAKYVLGADGARSGVRAAIGRKLEGTTSMHAWGVVDALITTDFPDYRNKATIQSNTAGNILWIPREGGFLTRLYVDLGEVTDENATEIRKTSLAHMEEICNEVLHPYSLQMKEVPWFSLYEVGHRVTDKFDNVPADETGKRTPRVFIAGDACHTHSAKAGQGMNVSMQDGWNLGWKLGYVLDGRSPESILDTYSKERQTIAVDLVTFDRTWSTIMAKKPEELTDPAERETFYVGTWEFPAGMKTQYPAAEMVGESTHQDLAKGFPIGKHFKSAHAMRVCDAKPVHIGHLHEADGRWRIYVFADSAAAAGKKSPTSDLAKWLETSPDSPLACQPKDGDLNAWFDVKVIYRQPWDEVEMAKVPPVFKPIMGPMCLQDWEQVFTVDPSDDIFVARQISDDGAIVVVRPDQYVAHVLPLTAHDELAAFFKPILLAK |
| MHBH3 | 3300045235\|Ga0485453_00791\|CDS\|Ga0485453_00791_7984_9894\|-\|7984:9894 | 45.7 | MQYHHDGYVGHDPRIREAAGTGIDRAEGLPDTMDVLIVGAGPAGIVQAAQLTDYPDVHTRIIERRPTRLAAGRADGLFPRSGETFQAFEFYDSIAQEAAHMREMSFWGPDPDNPTQVARGARAPDPPASVSEFPILTVNQARVIDYFAERAANGPARISIDYGYEFLDLTVHDSGEYPVEVRVRESDGTERTVRAKYVMGCDGARSRVRACIGAEVVSDSASQAWAVMDIMANTDFPDFRTRSGIQSDSNGTILLIPREGGFLTRFYVSLEAPTDESRERIMSTTVEEAIERANRILHPYSVDVRNVAWYSTYEVRHTVAQSFDDAAVEQVAGVTPRVFIAGDACHTHSAKAGQGMNVSIQDGWNLAWKIGQVLEGRSDESLLRTYHDERQDVAQKLIDYDKEWSAMMSKRPDEMEGPQEIVDFFVDTANFPGGFETRYQPSSIVGTAENQGLAAGFPIGMRFKSARVSRVADSLPVHVGHHFRADGRWRLYAFADGDGRAVADLAAWLEHAEDSPISLFTPAGADIDSVFDAKVIYQQGYRDVDFAAVPSLFRPKVGVYGVTDWEKVYATIPEDDFFDARGVDRSGALVIVRPDMYVAHVLPLSARAEITDFFAQSMVRQAATELVGAGVSRPTA |
| MHBH4 | 3300031110\|Ga0308165_1001428\|CDS\|Ga0308165_100142813\|+\|9603:11519 | 45.5 | MQIHYTGYESGDPRVKPAAGYGVDRPVELADEMDVLIVGGGPAATVVAAQLSRFPSINTYVIEQRPGRLEVGQADGIQARSMETFQTFGFAGEIVDEAYRNVETCFWKPDPADPTRIVRASRAPDDATGVSEFPHIYVNQSRVLDYFFRDAARAPGRVVPNYGIEFLDCTIHPDQDHPVEARVRYAVGPREGEVRSIRAKYLVGGDGARSRVRTSLGRRLRGQAALHAWGVMDVLLNTDFPDIQVKCLIQSQDAGSIITIPREGGYLVRFYVDLGDVDEAAVAGSVRATPVEDIIARANRILSPYVLDVKHVTWWSVYEVAHRIADGFDDVPEDEAGSRHPHAFIMGDACHTHSAKAGQGMNVSIQDGFNLGWKLAAVLERRSPESLLSTYAAERMVIAQNLIDFDKEWSDMLARSAGGLGDPDEVERYYLATIEFPAGFMTQYGPSMITSSDEHQSLAPGFPIGKRFKSAEAIRRADNRWRHIGHLHEADGRWRVYVFADSQAPTVSGSPVADFAWWWEADAASPQVRFTPPDLDPDAVFDTKVIYQQDYTEVEPANVPNAFKPVKGPYRLVDLNQIFSAGHGRDIFRDREVSREGAVVVVRPDQYVAGIFPLTAHDEIGAFFAGHLLEPISPADGR |
| MHBH5 | 3300010882\|Ga0123354_10014134\|CDS\|Ga0123354_1001413411\|-\|8843:10741 | 46.8 | MQFFLDGFRAGDPDVKPEAVRPTSDSDVVDVLVIGHGPAGALIAAQLSEFPGITTRLLERRETPMIQGQADGVNCRSIEMLNAFGLADRLNREGCFVNETAFWGANAAGTGIVRTGRVRDAEDGISEYRHVLLNQARLHEYLAEKMGNSPTRLQPEFGMEVIDLKLPDNPADLSQPDSVVVRQLNLPGQPETVIKAKYVIGCDGARSTVREALGLKLIGVPQFHGWGVLDVLATTDFPDIRRKVIIQAPGGGLAILPREGGYMTRLYVDLGDFEPGDNPRDRVSKDDIVAEAKRLFAPYTFEPKEIPWWSIYEVWQRVTERFDDVPDAERGKRSPRVFTAGDACHTHSAKIGGGMNLSMQDTFNLGWKLAAVLEGRSPVSLLDTYNDERMPLAHEFIEFDKVWAPMVDGKAPDGTAFSTEQLRNRFQEVEHYTAGVATHYPAALLVGKDTHQRLANGFTVGMRFHSAEVTRLADALRLHLGHTHEADGRWRLYAFGDAAPITSDSSRLRALCEWLVDSPDSPIRRYTAAGADIDAVFDVRGILQQSHNELEPEDLPSLLLPIKGSLGLNDYEKAFTRGVNDGDKDIFDVREIDRSQGALVIVRPDQYVANILPLDAYKELSDFFAGFMIPVK |
